# Sequence grammar underlying unfolding and phase separation of globular proteins

**DOI:** 10.1101/2021.08.20.457073

**Authors:** Kiersten M. Ruff, Yoon Hee Choi, Dezerae Cox, Angelique R. Ormsby, Yoochan Myung, David B. Ascher, Sheena E. Radford, Rohit V. Pappu, Danny M. Hatters

## Abstract

Aberrant phase separation of globular proteins is associated with many diseases. Here, we use a model protein system to understand how unfolded states of globular proteins drive phase separation and the formation of unfolded protein deposits (UPODs). For UPODs to form, the concentrations of unfolded molecules must be above a threshold value. Additionally, unfolded molecules must possess appropriate sequence grammars to drive phase separation. While UPODs recruit molecular chaperones, their compositional profiles are also influenced by synergistic physicochemical interactions governed by the sequence grammars of unfolded proteins and sequence features of cellular proteins. Overall, we find that the driving forces for phase separation and the compositional profiles of UPODs are governed by the sequence grammar of unfolded proteins. Our studies highlight the need for uncovering the sequence grammars of unfolded proteins that drive UPOD formation and lead to gain-of-function interactions whereby proteins are aberrantly recruited into UPODs.

**Highlights:**

- Unfolded states of globular proteins phase separate to form UPODs in cells
- The fraction of unfolded molecules and the sticker grammar govern phase separation
- Hydrophobic residues act as stickers that engage in intermolecular interactions
- Sticker grammar also influences gain-of-function recruitment into aberrant UPODs

## Introduction

Protein homeostasis (proteostasis) is achieved by protein quality control machineries that regulate protein production, folding, trafficking, and degradation (Balch et al., 2008; Powers et al., 2009). A major function of the proteostasis machinery is to facilitate the correct folding of globular proteins that have a stable fold (Bobori et al., 2017; Reinle et al., 2021; Sontag et al., 2017). We refer to these proteins as intrinsically foldable proteins (IFPs). In cells, IFPs have a broad range of protein folding stabilities (Leuenberger et al., 2017) that depend on protein sequence and fold type (**Figure 1A**). IFPs can be classified as being thermally unstable, U (bottom 10%), stable, S (top 10%) or of medium stability (remaining proteins). Analysis of the data of Leuenberger et al., shows that the sub-proteome comprising the least thermally stable IFPs (U) contain a higher fraction of disease related proteins compared to the sub-proteome comprising the most stable IFPs (S) (**Figure 1B**). Decreased stability leads to a higher proclivity for sampling non-native / unfolded states under folding conditions. Accordingly, IFPs that have a significant fraction of molecules in the unfolded state may leave the protein more susceptible to mutations that promote the formation of aberrant cellular deposits. Indeed, many IFPs with lower intrinsic folding stabilities have known disease mutations that further destabilize the folded state and / or promote concentration-dependent, aggregation-mediated phase separation that leads to the formation of aberrant cellular deposits (**Figure 1C** and **Table S1**)(Maier et al., 2009; Turner et al., 2005). For example, in the context of familial forms of amyotrophic lateral sclerosis (ALS), mutations to superoxide dismutase 1 (SOD1) affect the stability of the SOD1 dimer and promote the formation of protein deposits that accumulate through interactions among unfolded or partially unfolded monomeric states (Gomez and Germain, 2019; Meiering, 2008).

**Figure 1:**
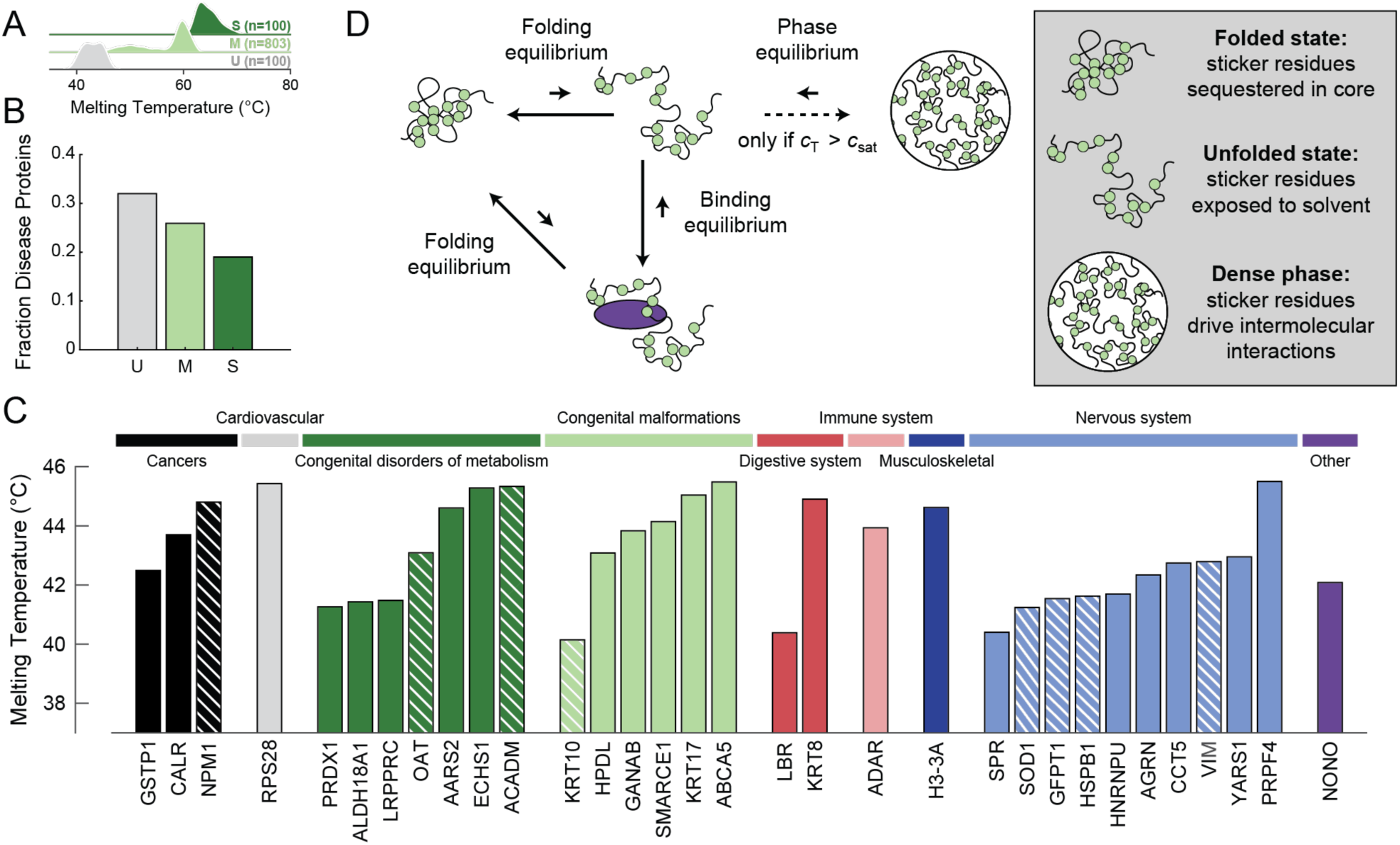
Normal cellular function requires proper balance of an interconnected equilibria triad of folding, binding to components of the quality control machinery, and phase separation. (A) Probability density estimates of the melting temperatures for human proteins in the unstable (U), medium stable (M), and stable (S) classes as defined by Leuenberger et al. and extracted from ProThermDB (**Table S1**) (Leuenberger et al., 2017; Nikam et al., 2020). In accordance with Leuenberger et al., we classify the bottom 10% of proteins in terms of melting temperature as unstable, the top 10% as stable, and remaining as medium stable. (B) The fraction of proteins in the U, M, and S classes that are associated with KEGG disease proteins (**Table S1**) (Kanehisa and Goto, 2000). (C) Melting temperatures of the 32 unstable human proteins associated with disease. Proteins are grouped by KEGG disease type. Stripes in the bars indicate that there is experimental evidence for disease associated mutations leading to aggregation-mediated phase separation (**Table S1**). (D) Schematic of the interconnected equilibria of folding, binding with protein quality control machinery, and phase separation. The green circles denote stickers, and the purple oval denotes chaperone binding. Here, *c*_tot_ denotes the total IFP concentration. When *c*_tot_ > *c*_sat_, the homogeneous well-mixed phase is saturated, and the system separates into two coexisting phases. See also **Figure S1** and **Table S1**.

IFPs in cells are defined by linked equilibria involving *folding* through intramolecular interactions, *binding* to components of the quality control machinery through specific heterotypic interactions, and *phase separation* through homotypic intermolecular interactions. The proposed triad of linked equilibria is inspired by findings from Frydman and coworkers (Kaganovich et al., 2008; Sontag et al., 2017), and it follows that the formation of deposits through phase separation of misfolded, partially unfolded, or unfolded proteins, driven by homotypic interactions, is part of the normal processing of unfolded proteins that is required for proteostasis (**Figure 1D**).

Our goal is to understand the principles that govern phase separation driven by homotypic interactions among unfolded or partially unfolded proteins. For IFPs that are two-state folders, the protein, under folding conditions, samples folded and unfolded states. Note that unfolded states sampled under folding conditions are distinct from the highly denatured state accessed in the presence of high concentrations of denaturants (Peran et al., 2019). The folding-unfolding equilibrium can be regulated by binding to components of the quality control machinery (Powers et al., 2009). For instance, chaperones generally bind to exposed hydrophobic patches of amino acids in the unfolded or partially folded states of IFPs to mediate the folding / re-folding process or to deliver IFPs for degradation. Accordingly, unfolded molecules can exist in two states: bound and unbound.

IFPs are also characterized by a phase equilibrium whereby they undergo concentration-dependent phase separation. These solvent quality (Crick et al., 2006; Pappu et al., 2008) mediated transitions are driven by homotypic interactions among unfolded molecules whereby, in its simplest form, a protein plus solvent system separates into a dilute, protein-deficient phase and a coexisting dense, protein-rich phase (Mathieu et al., 2020; Pappu et al., 2008; Posey et al., 2018a). Phase separation, which results from a combination of specific-and non-specific homotypic interactions, is a density transition, which we refer to as aggregation-mediated phase separation. Specific interactions arise from cohesive motifs known as stickers that form reversible physical crosslinks (Choi et al., 2019; Choi et al., 2020). For a given set of solution conditions, the strengths of driving forces for phase separation driven by homotypic interactions can be quantified by a saturation concentration, *c*_sat_, which is the threshold concentration of the protein above which it separates into coexisting dilute and dense phases (**Figure S1A**) (Wang et al., 2018). Thus, when the total concentration of protein is above *c*_sat_, a phase equilibrium exists between molecules in the dilute phase and molecules in the dense phase.

Although recent work has co-opted the term phase separation to refer to the coexistence of two liquid phases, it is worth noting that the formal definition of phase separation as a density transition does not impose any constraints on the material properties of coexisting phases. Indeed, the usage of saturation concentrations to quantify driving forces for forming protein-rich deposits via aggregation-mediated phase separation predates the current focus on liquid-liquid phase separation (LLPS) alone (Ciryam et al., 2017; Ciryam et al., 2013; Crick et al., 2006; Crick et al., 2013; Garai et al., 2008; Pappu et al., 2008). Phase separation can give rise to an assortment of coexisting phases, and the appropriate prefix such as liquid-liquid, liquid-solid, solid-solid etc., will depend on the material properties of the coexisting phases (**Figure S1B**).

For many IFPs, aberrant phase separation appears to be the result of concentration-dependent interactions among unfolded or misfolded proteins (Balchin et al., 2020; Clark, 2004; Hartl, 2016; McMillan et al., 2005; Ryno et al., 2013; Solomon et al., 2012; Song, 2018). For example, soluble wild-type (WT) SOD1 exists as a homodimer, stabilized by metal binding and an intra-subunit disulfide bond. Aberrant phase separation and formation of SOD1 deposits is driven by interactions among unfolded states of SOD1 (Nordlund et al., 2009). These results suggest several testable hypotheses. First, disease-related mutations likely restructure the triad of equilibria by increasing the concentration of unfolded molecules and thus decreasing the total protein concentration required to drive phase separation. Second, because phase separation appears to be driven primarily by interactions among unfolded molecules, the stickers that drive phase separation must be accessible in the unfolded state and drive homotypic interactions among unfolded molecules. Third, components of the protein quality control machinery can bind unfolded molecules and thereby weaken their ability to engage in homotypic interactions that lead to phase separation.

To test our hypotheses, we focused on answering the following questions: Are all unfolded states of IFPs equivalent as drivers of phase separation and the formation of aberrant, *de novo* unfolded protein deposits (UPODs) in cells, or must the unfolded states expose distinctive stickers that can drive phase separation? Can chaperones destabilize the formation of aberrant UPODs? Are all UPODs compositionally equivalent, or do different compositions of stickers recruit different proteins based on the physiochemical properties of the stickers? We answer these questions by utilizing the model protein barnase, a monomeric globular protein whose structure, stability and folding *in vitro* (Dalby et al., 1998; Matthews and Fersht, 1995) and *in vivo* (Wood et al., 2018) have been studied extensively.

Barnase is a bacterial ribonuclease. The catalytically inert H102A variant (referred to as the WT for the purposes of this study) is benign as far as mammalian cell physiology and viability are concerned (Wood et al., 2018). The steady-state population of molecules in folded vs. unfolded states is dictated by the free energy of unfolding: 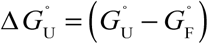. Here, 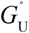 and 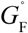 are the standard state free energies of the unfolded and folded states, respectively. The relative fraction of molecules in unfolded vs. folded states increases monotonically as 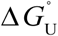 decreases in favor of the unfolded state. The fractions of molecules in folded vs. unfolded states are equal for 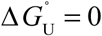. The equilibrium favors unfolded states if 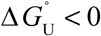. For proteins with large positive values of 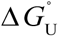 essentially ∼100% of the molecules will be in the folded state. Conversely, for proteins with large negative values of 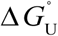 essentially ∼100% of the molecules will be in the unfolded state. Wood et al., measured the free energy of unfolding 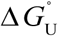 of a wide variety of barnase variants. These measurements provide a useful starting point for quantifying the relative, variant-specific concentrations of folded and unfolded molecules. Importantly, WT barnase, fused to mTFP1 at the N-terminus and Venus at the C-terminus, does not form visible deposits in mammalian cells (Wood et al., 2018). In contrast, variants for which 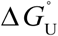 becomes less positive or even negative will have diminished stability. Increased access to unfolded states, through decreased stability, increases the concentration of unfolded proteins, and this leads to the formation of visible barnase deposits in mammalian cells. Unfolded states of barnase are also known to engage with components of the quality control machinery (Wood et al., 2018). Together these results imply that we can use barnase to interrogate the three-way interplay of protein stability, phase separation, and engagement with the quality control machinery.

Overall because barnase is orthogonal to the cellular system of interest, we can use it to investigate and uncover generic attributes of unfolded proteins that contribute to aggregation-mediated phase separation that leads to UPODs. Our findings, which include compositional profiling of UPODs formed by different barnase variants, allow us to understand how foreign UPODs engage with the cellular milieu. This is directly relevant for understanding how aberrant UPODs might engender toxicity. Finally, we provide evidence that the rules extracted using the model protein barnase are generalizable to disease associated human IFPs that undergo aberrant phase separation through homotypic interactions among unfolded molecules.

## Results

### Phase separation is driven by interactions among unfolded barnase molecules

We deployed an optoDroplet system to uncover the sequence grammar that underlies the phase separation of mutational variants of barnase molecules in live cells. The optoDroplet system was developed as a tool to study the phase separation of multivalent proteins in cells using a precise and controllable reaction triggered by blue light (Shin et al., 2017). The system involves a fusion of the protein of interest to the photoactivatable Cry2 domain (Hsu et al., 1996; Lin et al., 1998) and a fluorescent protein reporter (**Figure 2A**). Cry2 forms sub-microscopic oligomers upon blue light illumination. However, it does not drive phase separation on its own, even upon light activation (Lin et al., 1998). When fused to a domain that can undergo phase separation, the oligomerization of Cry2 reduces *c*_sat_ for the test protein allowing quantitative and inducible comparison of apparent *c*_sat_ values across different variants (Shin et al., 2017).

**Figure 2.**
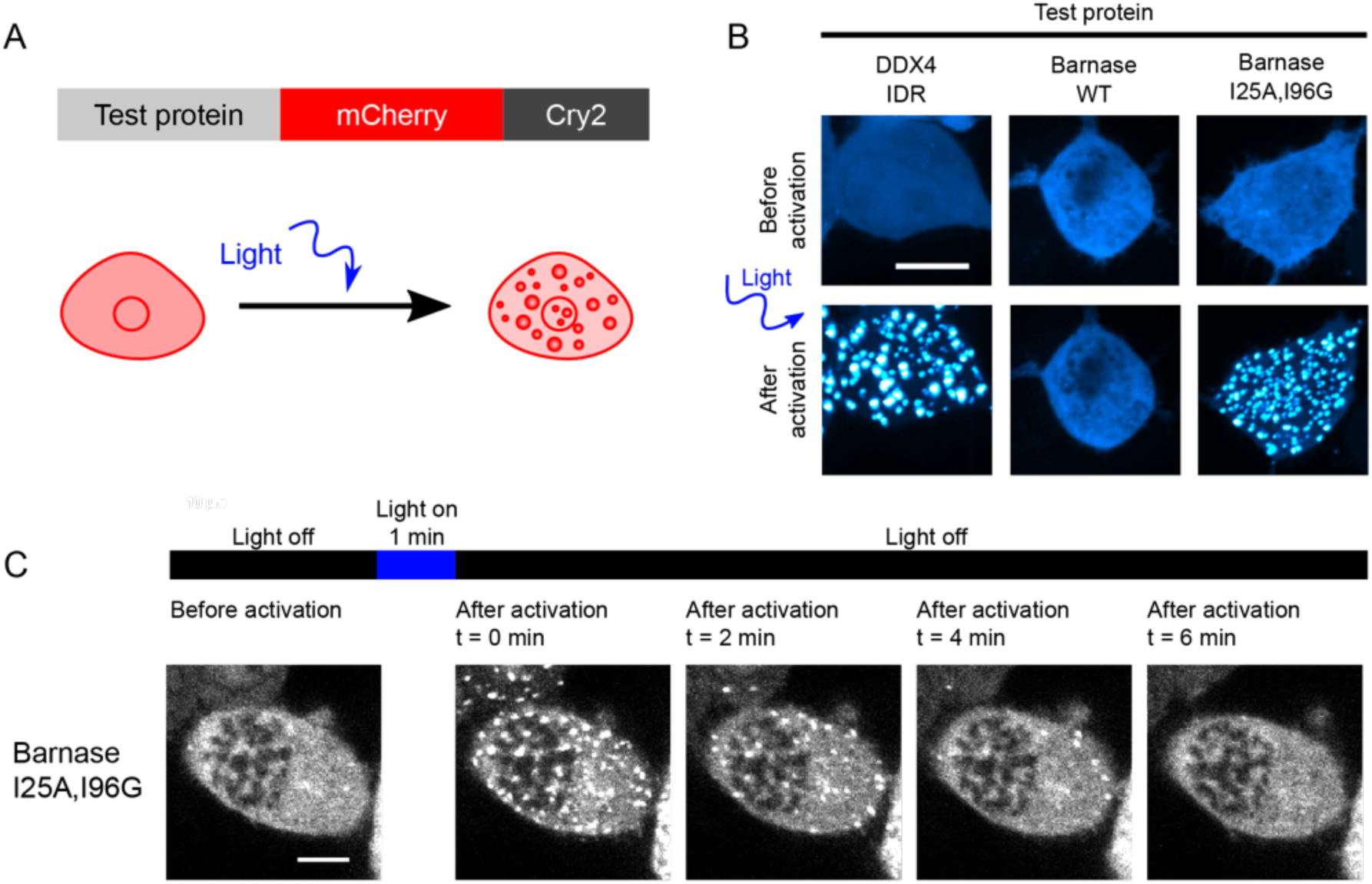
Phase separation is driven by interactions among unfolded barnase molecules. (A) Schematic of constructs used for the optoDroplet assay. (B) Representative confocal micrograph images of Neuro2a cells transfected with DDX4 IDR (positive control), WT barnase, and the destabilizing barnase variant (I25A, I96G) optoDroplet constructs before and after light activation. (C) Time lapsed confocal imaging of live Neuro2a cells expressing the I25A, I96G barnase optoDroplet construct. Scale bars in panels B and C correspond to 10 μm.

The intrinsically disordered region (IDR) of DDX4 was used as a positive control (**Figure 2B**) because it has been shown to undergo reversible phase separation in the optoDroplet setup (Brady et al., 2017; Nott et al., 2015; Shin et al., 2017). In contrast to the IDR of DDX4, WT barnase did not undergo phase separation when Cry2 was light activated (**Figure 2B**). However, the (I25A, I96G) double mutant, which has a finite probability of accessing unfolded states under physiological conditions, undergoes phase separation in a blue-light dependent manner (**Figure 2B** and **2C**). These results suggest that phase separation, driven by interactions among unfolded barnase molecules, can be assessed in a controlled manner without the confounding effects of slow kinetics that typically characterize the formation of UPODs in cells.

### A combination of protein destabilization and a distinct sequence grammar are required for UPOD formation via phase separation

We examined a range of mutations in barnase to titrate the impact of 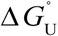 on phase separation. The values for these variants ranged from +18.7 kJ/mol (highly stable) to –0.8 kJ/mol (highly unstable) (Wood et al., 2018). Mutations with 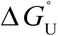 values above +13.0 kJ/mol were resistant to phase separation, whereas those with 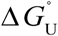 values below this threshold readily undergo phase separation (**Figure 3A**). If abundance of unfolded proteins, dictated by 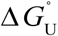, is the sole determinant of the driving forces for phase separation, then there should be a threshold concentration of unfolded proteins, *c*\*, above which the system separates into dilute and dense phases (**Figure 3B**). This concentration quantifies the saturation threshold of the unfolded species. The value of *c*\* defined as *c*\* = *p*_U_×*c*_sat_, is a product of the apparent saturation concentration, *c*_sat_, comprising folded and unfolded barnase molecules, and *p*_U_, which is the fraction of molecules in the unfolded state. Positive 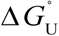 values imply that the fraction of unfolded proteins is less than the fraction of folded molecules, and thus a larger *c*_sat_ value is needed to reach *c*\*. Note that *c*\* ≈ *c*_sat_ as *p*_U_ approaches 1. Therefore, if phase separation is driven exclusively by the concentration of unfolded proteins, we expect that variants with the lowest fraction of unfolded proteins will have the highest *c*_sat_ values because *c*_sat_ = *c\**×(*p*_U_)^-1^.

**Figure 3.**
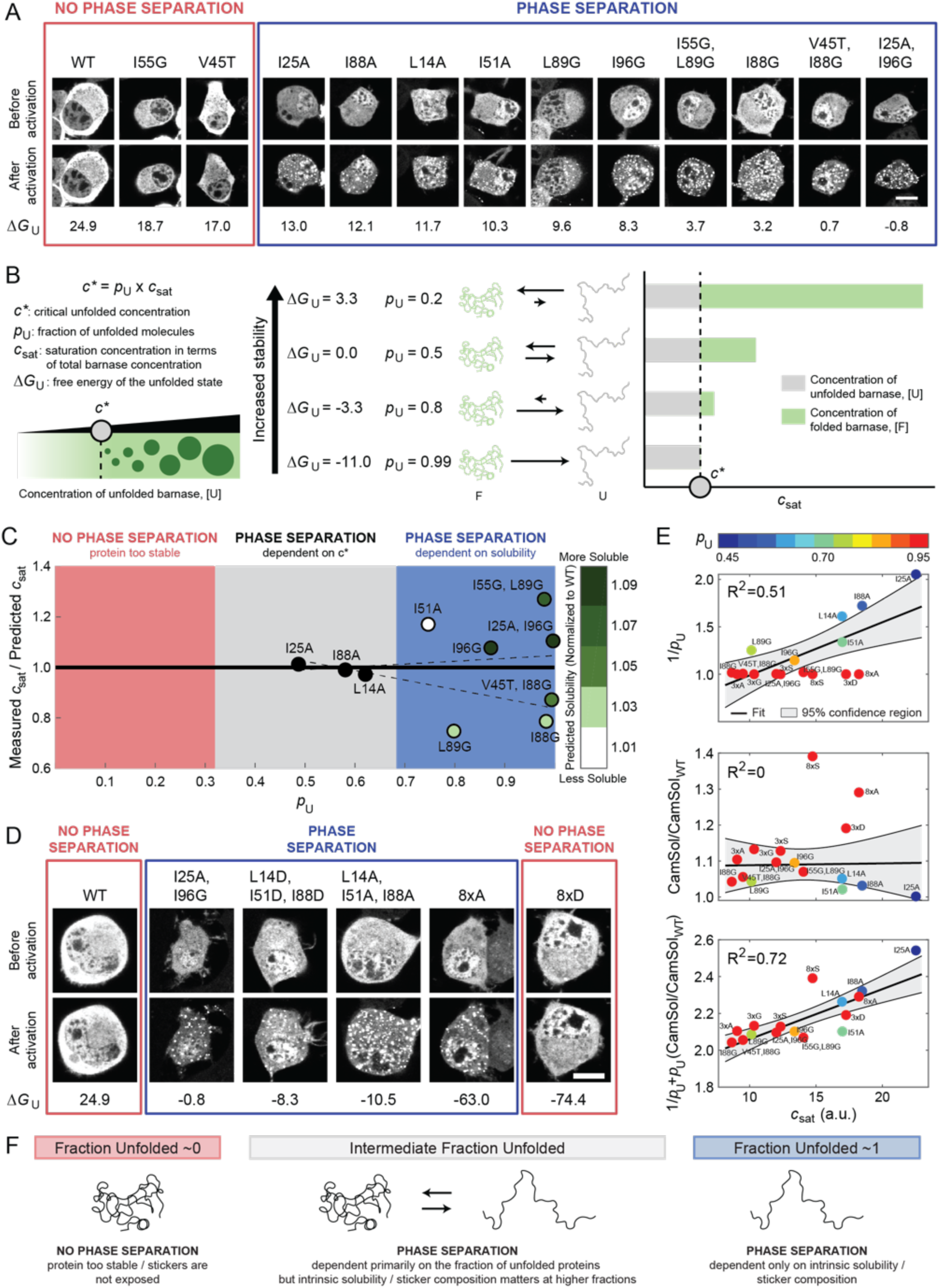
A combination of protein destabilization and a distinct sequence grammar are required for UPOD formation via phase separation. (A) Representative confocal micrograph images of Neuro2a cells transfected with the barnase-optoDroplet constructs as shown before and after light activation. Red box indicates constructs that do not undergo phase separation, whereas the blue box denotes constructs that do. Scale bar corresponds to 10 μm. 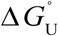 values are given in kJ/mol. (B) Model for how *c*_sat_ should change if stability, and thus a critical concentration of unfolded proteins, *c*\*, is all that matters for phase separation. (C) Measured versus predicted *c*_sat_ as a function of *p*_U_ (**Figures S2B** and **S2C, Table S2**). The predicted *c*_sat_ values were determined by globally fitting all barnase variants to *c*_sat_ = *c*\*/*p*_U_ for a constant *c*\* and an offset in Δ*G°*_U_ (**STAR Methods**). The best fit was for *c*\*=10.83 a.u. and 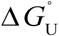 =-12.9 kJ/mol. Variants are colored by their normalized CamSol solubility score. Dashed lines denote the fitted confidence interval determined by 1000 bootstrapped trials of the variants, where the variants were picked based on the degree to which they modulated the hydrophobic and hydrophilic blobs from WT (**STAR Methods**) (D) Representative confocal micrograph images of Neuro2a cells transfected with the additional barnase-optoDroplet constructs with negative Δ*G°*_U_ values. Scale bar indicates 10 μm. Δ*G°*_U_ values are given in kJ/mol. (E) Comparison of *c*_sat_ and only stability (1/*p*_U_), only predicted solubility (CamSol/CamSol_WT_), or a combination of stability and solubility (1/*p*_U_+ *p*_U_(CamSol/CamSol_WT_)) using linear regression. Barnase variants are colored by their expected *p*_U_ given the offset in Δ*G°*_U_ of -12.9 kJ/mol. The 8xS variant is treated as an outlier in these analyses given the lack of cellular data at intermediate concentrations and thus the accuracy of the extracted *c*_sat_ is not clear (**Figure S2D**, grey box). (F) Summary of what features drive phase separation of IFPs. See also **Figure S2** and **Table S2**.

Previous studies have shown that many IFPs, including barnase, tend to have lower stabilities in cells than would be predicted based on estimates of 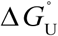 from *in vitro* measurements (Danielsson et al., 2015; Gnutt et al., 2019; Wood et al., 2018). Also, barnase has two appendages, Cry2 and mCherry, in the optoDroplet system. These are likely to alter the 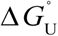 values when compared to estimates from *in vitro* measurements with untagged barnase molecules in dilute solutions. Accordingly, the analysis we deployed to estimate *c*\* uses a constant offset for 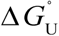 in relation to the 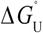 measured *in vitro* (see **STAR Methods**). For each barnase variant, we estimated *c*_sat_ using the dilute phase fluorescence intensity before and after light activation (**Figures S2A-C**). The lowest dilute phase fluorescence intensity at which we observe divergent behavior before and after light activation corresponds to the threshold concentration for the appearance of droplets (**STAR Methods**). The measured *c*_sat_ values are best described by a model where *c*\* ≈10.83 fluorescence intensity units (a.u.) that is based on an offset of -12.9 kJ/mol (−3.1 kcal/mol) for all 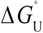 values (**Figure S2C**).

The analysis summarized in **Figure 3C** shows that the barnase variants fall into three categories. First, when the fraction of unfolded molecules is much less than the fraction of folded molecules, the concentration of unfolded molecules is too low and thus phase separation is not observed. Second, when the fraction of unfolded molecules is approximately equivalent to the fraction of folded molecules, phase separation is observed and the *c*_sat_ is accurately predicted by *c*\*. This implies that for variants in which ∼50% of the molecules are unfolded, the sequence of the variant has little impact on *c*_sat_. Third, when the fraction of unfolded molecules is high, *c*_sat_ is no longer accurately predicted by *c*\*. Instead, it appears that as *p*_U_ approaches 1, the underlying sequence grammar has a stronger influence on the driving forces for phase separation. Specifically, in this regime it is the intrinsic stickiness of the molecule that dictates the driving force for phase separation (Lang et al., 2015).

Mutations that destabilize the folded state of globular proteins often do so by weakening the hydrophobic core. Accordingly, if the residues that drive chain collapse and phase separation are equivalent (Bremer et al., 2022; Martin et al., 2020; Zeng et al., 2020), then destabilizing mutations would be expected to weaken the driving forces for phase separation of unfolded proteins. Therefore, we propose that even if the protein were completely unfolded (*p*_U_ ≈ 1), phase separation will only occur if the requisite sticker residues are present and accessible. To test our hypothesis, we used the CamSol method (Sormanni et al., 2015) to calculate relative solubilities normalized to that of WT barnase. The CamSol solubility score is calculated by considering local hydrophobicity, charge, and alpha-helix and beta-strand propensities, as well as hydrophobicity patterning and gatekeeping effects of charges. We find that all the mutational variants are predicted to be more soluble than WT. Furthermore, three of the four barnase variants that showed a higher measured *c*_sat_ compared to the predicted *c*_sat_ strongly increased the solubility compared to WT barnase (**Figure 3C**). These results suggest the following takeaways. First, when IFPs become primarily unfolded, the intrinsic solubility of the protein dictates the *c*_sat_. Second, the CamSol predictions suggest that interactions among hydrophobic residues of unfolded molecules are important for driving phase separation of IFPs.

To explore the importance of the requisite number of stickers being present and available for interactions among unfolded proteins, we introduced additional mutations into barnase that effectively ablated the folded state 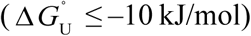. This enabled the quantification of driving forces for phase separation based solely on the properties of unfolded states (**Figures 3D, 3E**, and **S2D**). The mutations were chosen to alter the chemical environment of key bulky hydrophobic residues that would normally be in the core of the folded state. This includes triple mutations L14X, I51X, and I88X with X being A, G, S or D. These mutations are hereafter referred to as the 3×X variants. We also introduced octuple mutations L14X, L42X, I51X, L63X, I76X, I88X, L89X, I96X with X as A, S, or D, referred to as 8×X variants. In terms of hydrophobicity, the substitutions should follow the trend A > G > S > D (Kyte and Doolittle, 1982).

All variants except 8×D undergo phase separation in the concentration regimes that we explored (**Figures 3D, 3E**, and **S2D**). However, even though all variants were predicted to have a *c*_sat_ of ∼11 a.u. based on their *p*_U_, the measured *c*_sat_ values span a range from ∼9 to ∼18 a.u. (**Figure 3E** and **S2D**). These results suggest that not all unfolded states are equivalent as drivers of phase separation. Instead, sequence-specific stickers modulate the driving force for phase separation. Combining data for all the barnase variants, we find that neither *p*_U_ nor the intrinsic solubility of the unfolded state alone are suitable predictors of the measured *c*_sat_ values (**Figure 3E**, *R*^2^ = 0.51 and 0, respectively). However, consideration of both *p*_U_ and the intrinsic solubility of each variant correlates well with the measured *c*_sat_ values (**Figure 3E**, *R*^2^ = 0.72). This further suggests that the *c*_sat_ of IFPs with intermediate values of *p*_U_ will be dictated primarily by *p*_U_, whereas the *c*_sat_ of IFPs with *p*_U_∼1 will be dictated primarily by the intrinsic solubility of unfolded proteins (**Figure 3F**). Overall, the takeaway from these results is that phase separation requires that the unfolded state be favorably populated *and* that sticker-mediated interactions among unfolded molecules be minimally disrupted by mutations that destabilize the folded state.

### Phe and Tyr function as stickers that drive phase separation of unfolded barnase

To identify specific residues that function as stickers, we performed atomistic simulations of unfolded states of WT barnase. Residues predicted to be optimal stickers should have a higher probability of being in contact with other residues in the unfolded ensembles (Martin et al., 2020). We find that the residues with the highest mean contact probability are enriched in hydrophobic residues (**Figure 4A**). Of particular interest is the enrichment of Tyr and Phe given their roles as stickers that drive phase separation of intrinsically disordered prion-like low complexity domains (Bremer et al., 2022; Lin et al., 2017; Martin et al., 2020; Wang et al., 2018) and in forming the selectivity filter of nuclear pore complexes (Frey et al., 2006).

**Figure 4.**
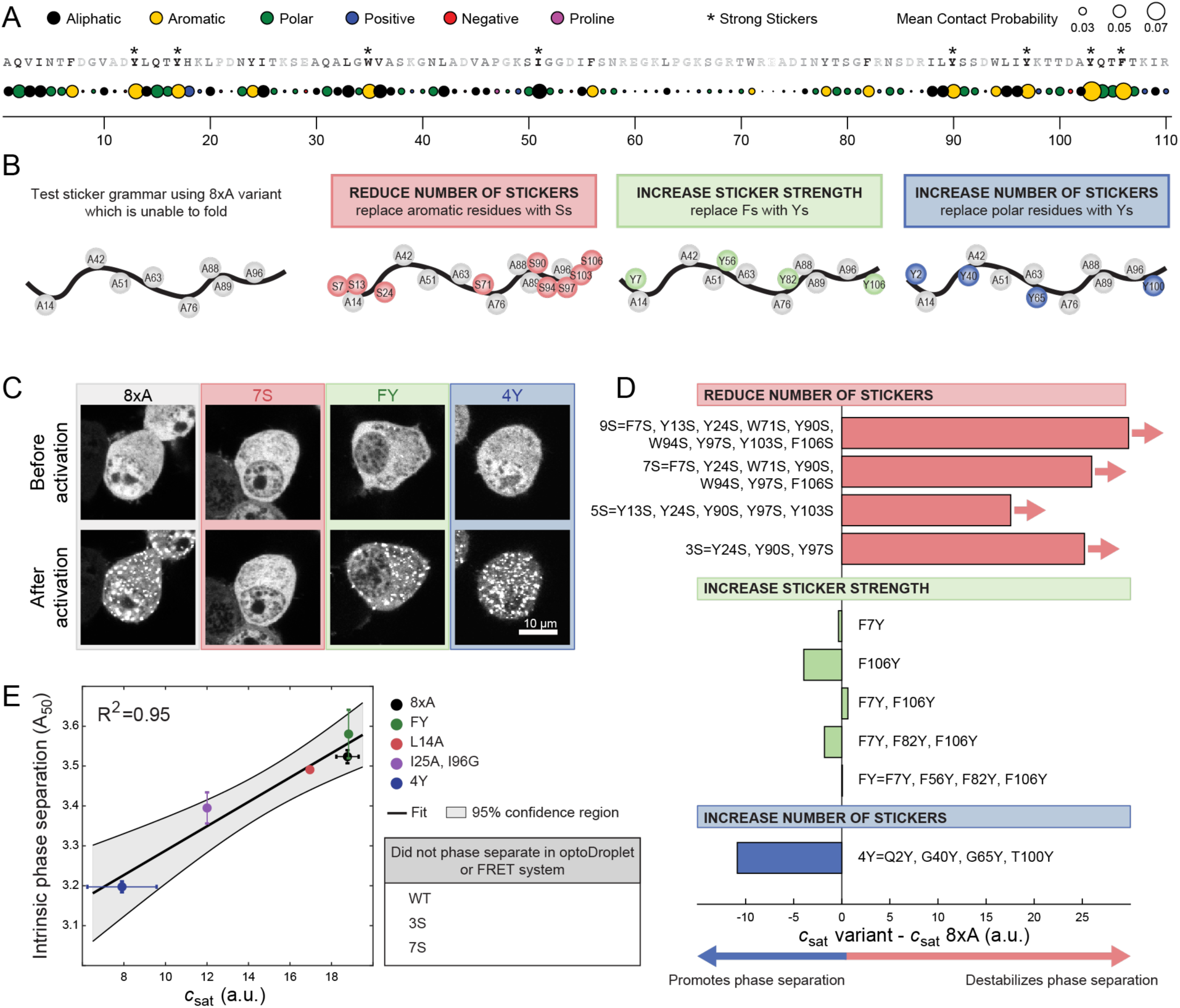
Phe and Tyr function as stickers that drive phase separation of unfolded barnase. (A) Mean contact probability for each residue quantified from atomistic simulations of unfolded states of WT barnase. For each residue, nearest and second nearest neighbour contacts are excluded from the calculation of the mean contact probability. The WT sequence is listed across the top and each residue is shaded based on its mean contact probability. Residues highlighted by * denote strong stickers. Strong stickers are those residues that have a mean contact order greater than the maximum mean contact order from a Flory Random Coil simulation (**STAR Methods**). (B) Schematic of constructs used to test sticker grammar. In all cases, the base construct is 8xA which has eight hydrophobic residues mutated to A (grey circles). Additional mutations used to test sticker grammar are shown in colored circles. Letters denote the residue the position is mutated to. (C) Representative confocal micrograph images of Neuro2a cells transfected with the sticker barnase-optoDroplet variant constructs. (D) Comparison of the *c*_sat_ values of each sticker barnase-optoDroplet variant construct with the *c*_sat_ of the 8xA construct, in arbitrary units. Bars with arrows indicate that a *c*_sat_ value could not be extracted for these constructs and must be at least above the value of the bar. (E) Comparison of the intrinsic phase separation of barnase as fusions to fluorescent proteins mTFP1 and Venus, using the A_50_ analysis, to *c*_sat_ values of barnase in the optoDroplet format (*R*^2^ = 0.95 for linear regression). Error bars indicate standard deviations. Also, see **Figure S3** and **Table S3**.

We tested the importance of aromatic residues as stickers for driving UPOD formation. To avoid confounding factors arising from the folded state, we introduced mutations into the 8×A variant, which is completely unfolded but still drives phase separation (**Figure 3D** and **3E**). Three categories of mutations were examined (**Figure 4B**). First, was the replacement of aromatic residues with Ser, which was predicted to reduce the number of stickers (Bremer et al., 2022). Second, was the replacement of Phe with Tyr, which was predicted to increase the sticker strength (Bremer et al., 2022). Third, was the replacement of polar residues with Tyr, which was predicted to increase the number of stickers (Bremer et al., 2022; Martin et al., 2020). The effects of these mutations were assessed using the optoDroplet assay (**Figures 4C, 4D and S3**). These measurements showed that decreasing the number of stickers weakened the driving forces for phase separation, whereas mutations of polar residues to Tyr enhanced the driving forces for phase separation. Substituting one or more Phe residues with Tyr had minimal impact on *c*_sat_. This suggests that Phe and Tyr have equivalent efficacy as stickers when phase separation is driven by interactions among unfolded barnase molecules.

### Interactions that drive phase separation of unfolded states have an equivalent impact on deposit formation

As noted in the introduction, phase separation is a density transition, and this is true irrespective of the material properties of the coexisting phases. UPODs typically form over long timescales and are not known to show the rapid reversible formation and dissolution that has been ascribed to liquid phases. If phase separation is a generic density transition, then we expect that the apparent *c*_sat_ values extracted using the optoDroplet assay should be equivalent to threshold concentrations extracted using an orthogonal assay that probes the formation of protein deposits in cells. We examined deposit formation using a FRET assay involving the barnase constructs fused to mTFP1 and Venus, where acceptor (Venus) fluorescence provides a readout on the local assembly of barnase molecules (Wood et al., 2018). Specifically, we derived estimates of the concentration of barnase in cells at which 50% of the cells contain deposits (A_50_ value). The lower this concentration, the stronger the driving force for deposit formation. We found a strong positive correlation between the A_50_ and *c*_sat_ values (*R*^2^ = 0.95 for linear regression) (**Figure 4E**). These measurements demonstrate the equivalence of driving forces for deposit formation and droplet formation in the optoDroplet assay. Furthermore, variants that did not form deposits also did not form droplets. Therefore, phase separation is a density transition, and this is true irrespective of the material properties of the coexisting phases.

### Molecular chaperones suppress phase separation of unfolded barnase molecules

Proteostasis is achieved through the collective actions of chaperones that bind to unfolded and / or partially unfolded states of proteins and mediate their processing for folding or delivery to other cellular machinery (Hartl, 2016). Here, we sought to explore how chaperones influence phase separation driven by interactions among unfolded barnase molecules.

Upon phase separation, unfolded proteins coexist in dilute and dense phases, the latter defined by protein deposits. Components of the chaperone system can bind to unfolded barnase either in the dense (**Figure S4**) or dilute phase (Wood et al., 2018). If binding to unfolded proteins in the dilute phase is stronger than in the dense phase, then *c*_sat_ in the presence of the chaperone, designated as 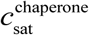, will be greater than *c*_sat_ in the absence of the chaperone. Conversely, if binding to unfolded proteins in the dilute phase is weaker than in the dense phase, then 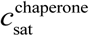 will be lower than *c*_sat_ in the absence of the chaperone. Preferential binding, which would be true of chaperones that function independently of ATP hydrolysis, is referred to as polyphasic linkage (Ruff et al., 2021b; Wyman and Gill, 1980). Upon binding of unfolded proteins by chaperones there should be three states of barnase in the dilute phase. These are folded barnase, unfolded barnase, and unfolded barnase bound to chaperones (**Figure 5A**). Previous studies showed that members of the Hsp70 and Hsp40 families can bind to barnase in the dilute phase and suppress UPOD formation (Wood et al., 2018). Accordingly, we proposed that while the total concentration of unfolded proteins (free + bound) would be higher in the presence of chaperones, the fraction of molecules capable of phase separation should be lowered (**Figure 5A**).

**Figure 5.**
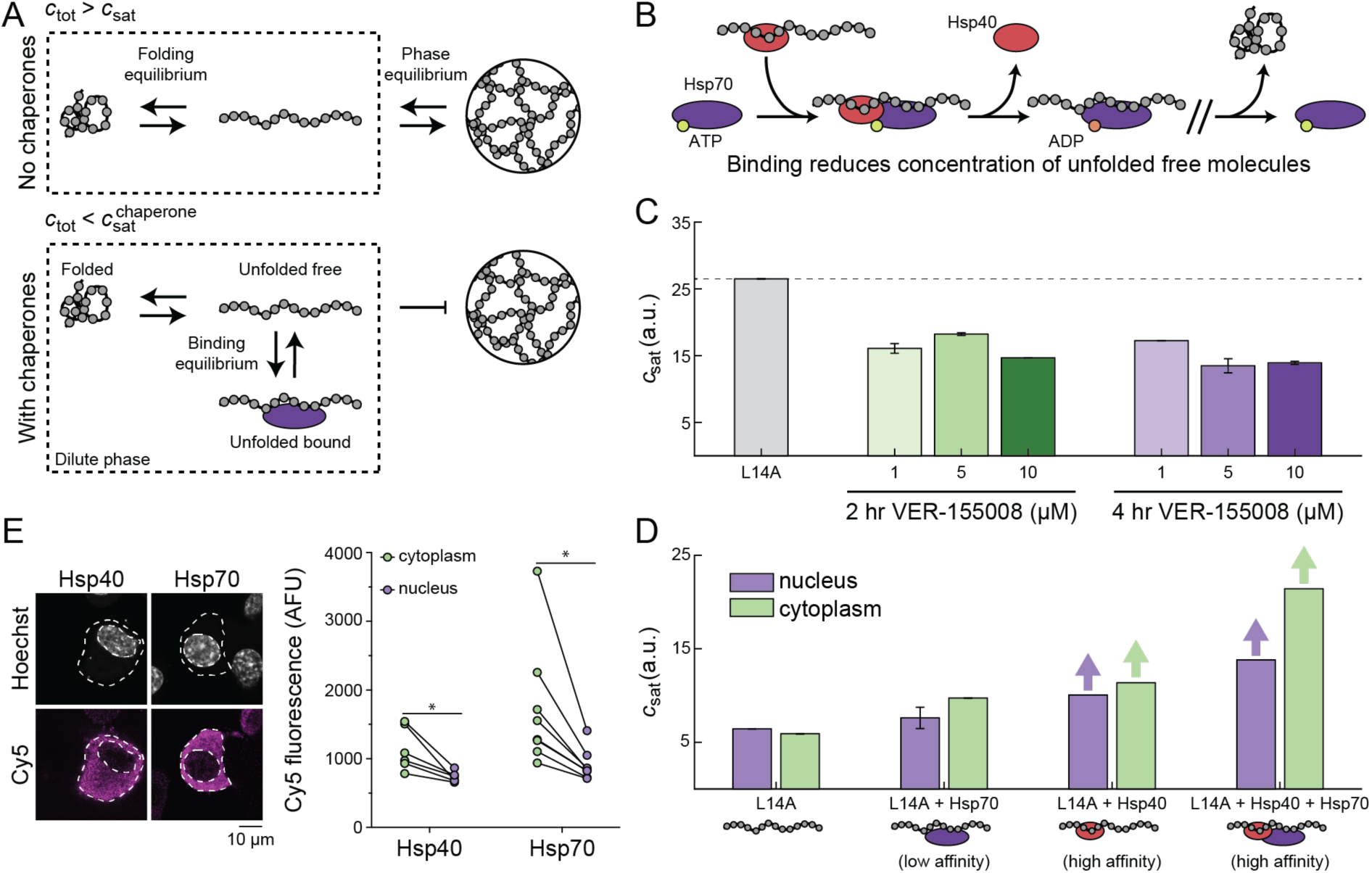
Molecular chaperones suppress phase separation of unfolded barnase. (A) In the absence of chaperones, the dilute phase consists of two dominant states, folded and unfolded barnase. There exists a phase equilibrium when the total concentration of barnase, *c*_tot_, is greater than *c*_sat_ and thus phase separation occurs. In the presence of chaperones, barnase in the dilute phase consists of three dominant states: folded, unfolded free, and unfolded bound to chaperones. At the same total concentration of barnase as in the absence of chaperones, barnase cannot phase separate because *c*_tot_ is less than the saturation concentration needed in the presence of chaperones, 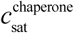. This results from the fact that chaperone binding reduces the concentration of free unfolded barnase. (B) Basic model for chaperone function. Hsp40 binds the unfolded molecule and forms a ternary complex with Hsp70 in the ATP-bound state. ATP hydrolysis leads to the release of Hsp40 and the formation of a high affinity complex between Hsp70 and the unfolded substrates. (C) *c*_sat_ for the L14A barnase-optoDroplet variant construct transiently transfected in Neuro2A cells in the absence or presence of different dosages of the Hsp70 inhibitor VER-155008. Dashed line corresponds to the *c*_sat_ of L14A in the absence of the inhibitor. Error bars denote the standard deviation from 50 bootstrapped trials. (D) *c*_sat_ for the L14A barnase-optoDroplet variant construct in the absence or presence of overexpressed chaperones. Bars with arrows indicate a *c*_sat_ value could not be extracted for these systems and must be at least above the value of the bar. Error bars denote the standard deviation from 50 bootstrapped trials. (E) Confocal images of Neuro2a cells transfected with V5-tagged DNAJB1 (Hsp40) or HSPA1A (Hsp70). Cells were stained by immunofluorescence for the V5-tag (Cy5) and the nucleus was stained with Hoechst 33342. Graphs show quantitation of immunofluorescence. Data shown as paired samples from individual cells. Paired t-test results shown; * p<0.05. See also **Figure S4** and **Table S4**.

The canonical model is that Hsp40 binds substrates, and then forms a ternary complex with Hsp70 in the ATP-bound state (Alderson et al., 2016; Jiang et al., 2019) (**Figure 5B**). ATP hydrolysis correlates with release of Hsp40 and the formation of a high affinity complex between substrate and Hsp70. If unfolded barnase molecules are a target of the Hsp40 / Hsp70 system, we expect that inhibiting Hsp70 should promote phase separation and decrease the *c*_sat_ of barnase molecules. Indeed, we find that inhibiting Hsp70 by the compound VER-155008 (IC50 = 0.5 μM) causes a lowering of the optoDroplet estimated *c*_sat_ of variant L14A (**Figure 5C**). Inhibition of Hsp70 by the small molecule appears to increase the pool of unbound unfolded proteins, thereby lowering *c*_sat_.

To further assess the impact of chaperones on the phase separation of destabilized barnase molecules, we co-expressed the optoDroplet construct containing the L14A barnase variant with the Hsp70 protein HSPA1A and / or its cofactor, the Hsp40 protein DNAJB1. Because the ternary complex is needed for Hsp70 to stimulate ATP hydrolysis and form a high affinity complex with unfolded proteins, overexpression of Hsp70 alone should result in fewer unfolded proteins being bound by chaperones when compared to Hsp40 alone or Hsp70 overexpressed with Hsp40. We found that overexpressing chaperones suppressed droplet formation of L14A barnase (**Figure 5D**). Overexpression of Hsp70 alone had the smallest effect on suppression of droplet formation, whereas droplets were not observed when Hsp40 was overexpressed, or when Hsp70 was jointly overexpressed with Hsp40. Additionally, we found that suppression of droplet formation was more pronounced in the cytoplasm than in the nucleus. This finding is consistent with overexpressed Hsp70 and Hsp40 accumulating predominantly in the cytoplasm, as confirmed by immunostaining (**Figure 5E**).

### UPODs can sequester and enrich cellular proteins through interactions that are governed by physical chemistry alone

Thus far, we have focused on uncovering the features that drive phase separation of unfolded molecules, and how phase separation can be modulated by molecular chaperones. To understand the physiological consequences of phase separation driven by unfolded molecules we need to understand how UPODs engage with the surrounding cellular milieu. It has been hypothesized that aberrant phase separation may disrupt cellular functions by at least two mechanisms. First, aberrant phase separation may recruit proteostasis machinery and thus modulate the balance of homeostasis (Hipp et al., 2014; Stefani and Dobson, 2003). Second, aberrant phase separation may lead to the sequestration and loss of function of unrelated proteins (Olzscha et al., 2011; Wear et al., 2015). To test for both possibilities, we undertook a compositional profiling of UPODs formed by different barnase variants.

We used a proteomics-based strategy to profile the protein compositions of UPODs formed by eight different barnase variants: WT, L14A, Ex4 (I25A, I96G), 8×A, 3S, 9S, FY, and 4Y (**Figure 6A-B**). Briefly, HEK293T cells were transfected with each of the eight barnase variants. Digitonin lysis was then performed, and diffuse proteins were removed. The compositions of the remaining insoluble material, which retained the variant-specific barnase UPODs, were quantified using mass spectrometry (**STAR Methods**). The barnase variants we chose have a range of stabilities, *c*_sat_ values, and sticker compositions (**Figure 6B**). As expected, barnase was the most abundant protein in the insoluble fraction for all barnase variants, except WT, in accordance with unfolded molecules forming UPODs (**Figure S5A**). Additionally, the abundance of barnase was highest for the variants that underwent phase separation (**Figure 6C**).

**Figure 6:**
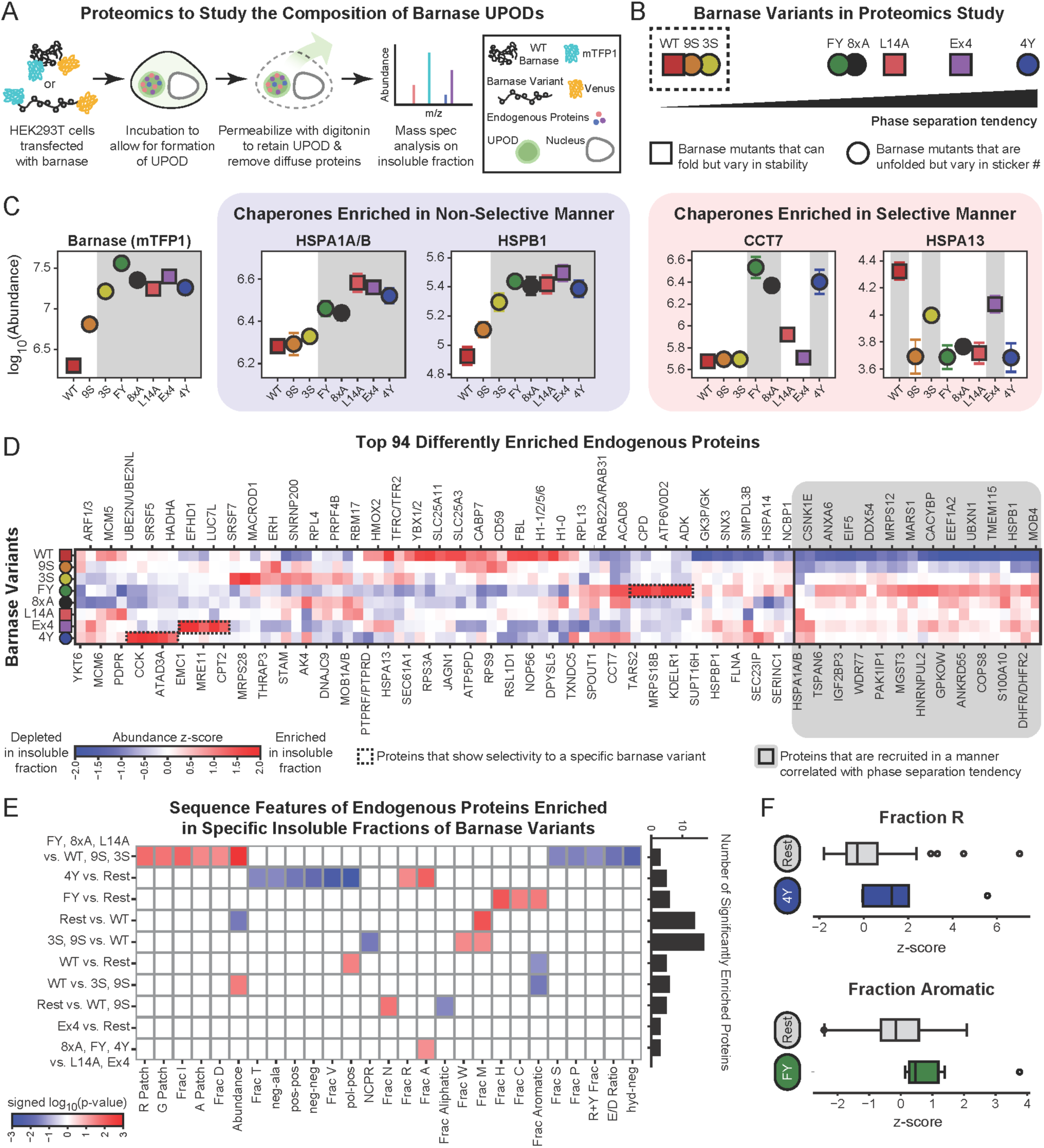
UPODs can sequester and enrich cellular proteins through interactions that are governed by physical chemistry alone. (A) Schematic of the proteomics workflow to extract the compositional profiles of specific barnase variant UPODs through analysis of the insoluble fractions of cells. (B) Description of the barnase variants used for the proteomics study. Barnase variants vary in phase separation tendency (*c*_sat_), stability (Δ*G°*_U_), and sticker composition. Variants within the dashed box correspond to variants that were not found to phase separate at the concentrations tested. (C) Abundance of barnase (mTFP1) and four representative chaperones in barnase specific insoluble fractions. Barnase variants are sorted based on their phase separation tendency. Selectivity refers to recruitment not correlated with the phase separation tendency of the barnase variants. Shaded gray regions denote which barnase UPODs the given protein is significantly enriched in as determined by the Fisher’s LSD test following an ANOVA test. Error bars denote the standard error of the mean of four replicates. (D) Smoothed abundance z-score matrix for the top 94 differently enriched proteins in the insoluble fractions (Cox et al., 2022). Here, the z-score is calculated using the mean and standard deviation of all replicas and all barnase variants for a given endogenous protein. Proteins are hierarchically clustered using the Euclidean distance and Ward linkage method. The 24 proteins highlighted in grey are recruited in a manner correlated with the phase separation tendency of the barnase variant. (E) Significant sequence features in different protein sets. The given set of proteins are significantly enriched in the insoluble fractions of barnase variants to the left of “vs.” compared to the barnase variants to the right of “vs.” (**STAR Methods**). Here, “Rest” refers to all remaining barnase variants. Features come in three types: patterning (i.e., pos-pos), composition (i.e., Frac R), or abundance (**STAR Methods**). Blue boxes denote either compositional features / abundance that are significantly depleted or patterning features that are well-mixed in the given protein set. Red boxes denote either compositional features / abundance that are significantly enriched or patterning features that are blocky in the given protein set. Significance is determined by using the two-sample Kolmogorov-Smirnov test on the z-score feature distribution of the given protein set compared to the z-score feature distribution of the remaining top 94 proteins. Bar chart shows the number of significantly enriched proteins in each set. (F) Boxplots of the z-scores of the fraction of Arg in proteins significantly enriched in the 4Y insoluble fraction vs. Rest and fraction of aromatics in proteins significantly enriched in the FY insoluble fraction vs. Rest. Grey boxplots denote the z-scores of the remaining top 94 differently enriched proteins in each case. See also **Figure S5** and **Table S5**.

Several of the topmost abundant proteins were found to be chaperones (**Figure S5A**). To understand how UPODs engage with the surrounding cellular milieu, we examined the abundance of different chaperones in the barnase specific insoluble fractions. We found that certain chaperones were enriched in a manner that was correlated with the phase separation tendency of the barnase variants (**Figure 6C**). These chaperones included HSPA1A/B and HSPB1. This suggests that certain chaperones are recruited to UPODs in a way that is largely non-selective with respect to the sticker composition of the barnase variant. In this case, all barnase variants are likely to be equivalent substrates and the chaperones act in a non-selective manner to maintain the proper balance of folding, binding, and phase equilibria.

In contrast, we found that other chaperones were enriched in the barnase specific insoluble fractions in a selective manner that was not correlated with the phase separation tendency of the barnase variants (**Figure 6C**). These chaperones included CCT7 and HSPA13. Specifically, CCT7 was enriched in 8×A, FY, and 4Y UPODs. These variants are all completely unfolded and have exposed aromatic residues. The combination of these features makes 8×A, FY, and 4Y distinct from the other barnase variants and suggests that the accessibility of stickers make them specific substrates for CCT7.

CCT7 is a subunit of the chaperonin-containing T-complex (TRiC) (Spiess et al., 2004). TRiC is composed of eight paralogous subunits, CCT1-8. All CCT subunits use the same region of the apical domain to interact with substrates (Spiess et al., 2006). For each individual subunit, this region is highly conserved across orthologous subunits (Joachimiak et al., 2014). However, each paralogous subunit has its own sequence composition preferences. These preferences lead to substrate specificity among the CCT subunits. Does the sequence composition of CCT7 explain why it targets the 8×A, FY, and 4Y variants? Indeed, the apical domain of CCT7 has the highest fraction of aromatic residues when compared to the other seven subunits (**Figure S5C**). Additionally, the aromatic residues are localized to the region of the apical domain important for substrate specificity (**Figure S5D**). Thus, it appears that the increased accessibility of aromatic residues in 8×A, FY, and 4Y and the increased aromatic fraction in the substrate recognition region of CCT7 makes these barnase variants specific substrates to CCT7 through interactions involving aromatic residues. This is consistent with previous results showing that mutating a single Trp in the β-isoform of the thromboxane A_2_ receptor reduces its interaction with CCT7 (Génier et al., 2016). Overall, our results suggest that certain components of the proteostatic machinery are generically recruited to UPODs to resolve them. However, other chaperones show selectivity based on the barnase variant. This selectivity suggests that properties of the individual barnase variant will determine whether it is a substrate of the chaperone.

We next asked whether other proteins enriched in the insoluble fractions also showed generic vs. selective recruitment. **Figure 6D** shows the abundance of the top 94 differently enriched endogenous proteins identified by a one-way ANOVA (**STAR Methods**). Of the 94 proteins, 24 were enriched in the insoluble fractions in a manner that correlated with the underlying phase separation tendency of the barnase variant (**Figure 6D**, grey solid box). The remaining 70 proteins showed different types of selectivity. Interestingly, there were subsets of endogenous proteins that were selectively enriched in insoluble fractions of specific barnase variants (**Figure 6D**, dashed boxes).

To identify and classify the distinct set of proteins associated with UPODs driven by different barnase variants, we identified proteins that were significantly enriched in a specific barnase insoluble fraction or a set of barnase insoluble fractions. For this, we used a *post hoc* Fisher least significant difference (LSD) test following an ANOVA test. For the identified sets of proteins, we did not find statistically significant results in GO cellular component, GO molecular function, or GO biological process, when the entire identified protein set was used as a reference. This result suggests that barnase specific recruitment is not due to shared cellular functions, processes, or localization among the enriched proteins (**Figure S5E**).

We hypothesized that UPOD specific recruitment might be due to physiochemical properties of the proteins such as complementary interactions with specific stickers that make up each of the barnase variants. To test for this possibility, we extracted ∼90 unique sequence features and compared the distribution of these features in each enriched set to the top 94 proteins using the two-sample Kolmogorov-Smirnov test (**Figure 6E**). We found that recruitment to barnase specific insoluble fractions depends on the underlying grammar of the specific barnase variant. For example, proteins that are only enriched in 4Y UPODs show a higher fraction of Arg residues, and these residues are dispersed uniformly along the linear sequence (**Figure 6E and 6F**). This is consistent with results showing that the numbers of Tyr and Arg residues jointly contribute to the co-condensation in FET family proteins (Wang et al., 2018). The additional Tyr residues in 4Y might explain why UPODs formed by this variant are enriched in Arg-rich proteins when compared to 8×A and FY. Additionally, for proteins that are only enriched in UPODs formed by the FY variant, we observe an enrichment of proteins with higher fractions of aromatic residues (**Figure 6E and 6F**). This result is consistent with the fact that Tyr is a stronger sticker than Phe (Bremer et al., 2022).

Taken together, the implication is that UPODs can recruit and sequester cellular proteins through interactions that are governed by physical chemistry alone, without any regard to overlapping or synergistic biological functions. This finding suggests that UPODs might enable gain-of-function interactions that deplete cells of key proteins. It therefore follows that protein-rich deposits that form in the context of disease may have idiopathic effects on toxicity through dysfunction caused by grammar-specific gains-of-function that are manifest in the form of UPOD-specific compositions.

### The sequence grammar that drives phase separation of unfolded states appears to be similar between barnase and disease associated IFPs, including SOD1

We have shown that phase separation of mutational variants of barnase requires that the concentration of unfolded molecules cross the *c*\* threshold and that these molecules have the requisite types and numbers of stickers to drive homotypic interactions among unfolded molecules. The relevant stickers are hydrophobic residues that become accessible for intermolecular interactions in the unfolded state. We focused our studies on barnase as an orthogonal model protein in mammalian cells. Do the rules gleaned from studies of barnase transfer to endogenous IFPs from human cells? To answer this question, we performed atomistic simulations of unfolded states for six different unstable IFPs from the human proteome (**Figure 1C**). We find that all six proteins feature stickers in the unfolded state that are either aliphatic and / or aromatic residues (**Figure 7A**). These residues account for a large fraction of the total mean contact probability for each protein and this fraction is larger than what would be expected based purely on their numbers in the sequences. In contrast, while polar residues also account for a large fraction of the total mean contact probability, this fraction is consistent with the number of polar residues in the sequence – a feature that is consistent with polar residues often being spacers instead of drivers of phase separation.

**Figure 7:**
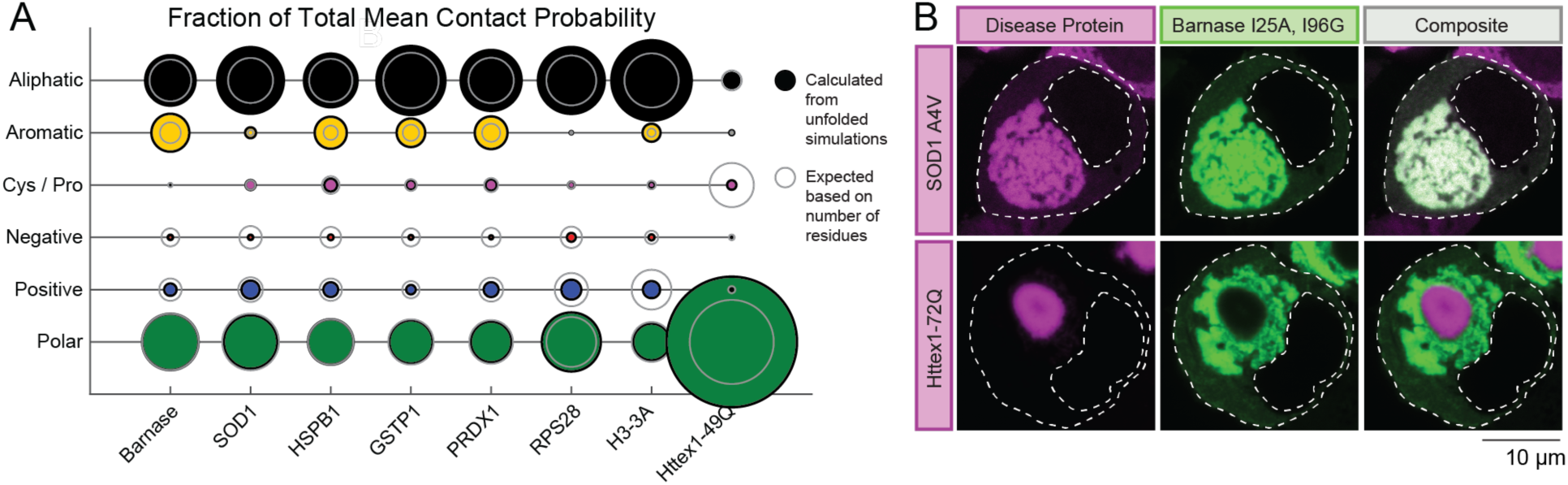
The sequence grammar that drives phase separation of unfolded states appears to be similar between barnase and disease associated IFPs, including SOD1. (A) Fraction of total mean contact probability per residue type calculated from atomistic simulations of the unfolded state (**STAR Methods**). Residue types with a high fraction of total mean contact probability that is greater than expected are likely going to be the predominant stickers. (B) The fluorescence micrographs show deposits formed by a destabilized variant of barnase (I25A, I96G) flanked with fluorescent proteins (mTFP1 and Venus) (Wood et al., 2018) along with deposits formed by mutant SOD1 (SOD1 A4V) or mutant Httex1 containing a glutamine tract of 72 residues (Httex1-72Q) fused to mCherry. The constructs were co-transfected in HEK293T cells. The outlines of the cells and nuclei are shown with dashed lines

We also examined which residues act as stickers in polyglutamine (polyQ)-expanded Huntingtin exon 1 (Httex1), a protein that is disordered and forms amyloid-like solids in cells (Bäuerlein et al., 2017). In contrast to the IFPs, polar residues dominate the fraction of total mean contact probability in Httex1 with an expanded polyglutamine tract of 49 residues. This fraction is greater than expected and the result is consistent with studies showing that the phase behavior of Httex1 is driven mainly by amide-amide interactions involving the polyQ domain (Crick et al., 2013; Posey et al., 2018b). These interactions are distinct from hydrophobic interactions anticipated to be responsible for driving phase separation of the IFPs studied here.

To test whether IFPs have a similar sticker grammar that is distinct from Httex1, as predicted from simulations, we assessed the colocalization of UPODs formed by the destabilized double mutant of barnase (I25A, I96G, 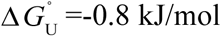) with a destabilizing mutant of SOD1 (A4V) and Httex1 with a glutamine tract of 72 residues (Httex1-72Q). Colocalization would imply that phase separation is governed by similar driving forces, i.e., a similar sticker grammar. When co-expressed in HEK293T cells, barnase I25A, I96G formed deposits that colocalized with those of SOD1 A4V (**Figure 7B**). We interpret these results to imply that phase separation of unfolded barnase and SOD1 are driven by similar interactions. In contrast, the barnase I25A, I96G UPODs did not co-localize with Httex1-72Q deposits (**Figure 7B**). Previous work has also shown SOD1 and Httex1 deposits do not co-localize (Farrawell et al., 2015; Polling et al., 2014). This lack of colocalization supports the hypothesis that distinct interactions underlie the phase behavior of SOD1 and barnase variants when compared to Httex1. Furthermore, these results suggest that interactions among hydrophobic residues in the unfolded states will likely drive phase separation for many IFPs.

## Discussion

In this work, we showed that interactions among unfolded states of IFPs drive intracellular phase separation leading to the formation of *de novo* deposits (UPODs). The formation of UPODs is influenced by two features. Phase separation is thermodynamically favored if the protein has a large enough concentration of unfolded proteins *and* has the requisite valence (number) and strength of stickers. The concentration of unfolded proteins is dictated by the free energy of unfolding, 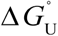, whereas the sticker valence and strength are dictated by the composition, accessibility, and sequence contexts of stickers in the unfolded states.

The specific stickers for IFPs appear to be aliphatic and aromatic residues. Interestingly, aromatic residues in many intrinsically disordered domains also drive the formation of distinct biomolecular condensates (Frey et al., 2006; Martin et al., 2020). The computational approach we deployed here to identify stickers (**Figures 4A** and **7A**) is, in principle, deployable in conjunction with advances in machine learning (Russ et al., 2020) across the unfolded proteome to make quantitative predictions and identify residues that act as stickers to drive the formation of UPODs.

We showed that the driving forces for forming UPODs can be modulated by molecular chaperones. Specifically, we found that preferential binding of chaperones to unfolded proteins in the dilute phase leads to a destabilization of UPODs. Our findings suggest that the modulation of phase separation by preferential binding of chaperones to the dilute phase represents thermodynamic control through polyphasic linkage (Ruff et al., 2021a, b; Wyman and Gill, 1980), to the regulation of the concentrations of free unfolded proteins. It is worth noting that while the action of Hsp70 is likely to involve a combination of preferential binding and ATP hydrolysis, Hsp40 functions purely through preferential binding. Interestingly, overexpression of Hsp40 has a stronger effect than Hsp70 alone, and the combination of the two has the strongest inhibitory effect on phase separation.

Consistent with the idea that chaperone binding is important for the proper regulation of UPODs, we found that HSPA1A/B and CCT7 were among the most highly abundant proteins in barnase UPODs (**Figure S5A**). Unlike CCT7, HSPA1A/B, a Hsp70 protein, is recruited to barnase UPODs in a manner that correlates with the phase separation tendency of barnase. This suggests that the underlying sequence composition of the substrate has little effect on HSPA1A/B recruitment. However, Hsp70 proteins are often not the first chaperones to bind unfolded substrates. Instead, they are recruited through interactions with other chaperones, including Hsp40s and small heat shock proteins (sHsps) (Alderson et al., 2016; Veinger et al., 1998). Interestingly, we found that the sHsp HSPB1 is also recruited to barnase UPODs in a non-selective way. Like Hsp40s, HSPB1 is an ATP-independent chaperone and thus its binding to unfolded proteins and modulation of UPOD formation can also be described by polyphasic linkage (Jakob et al., 1993; Ruff et al., 2021a, b). Additionally, HSPB1 has been shown to be one of the most promiscuous sHsps, consistent with the lack of selectivity for distinct barnase UPODs (Mymrikov et al., 2017).

HSPB1 functions by co-assembling with substrates (Gonçalves et al., 2021; Żwirowski et al., 2017). Co-assembly allows for substrates to be held in a proper state needed for Hsp70 dependent disassembly and refolding. Additionally, during this process, Hsp70 and its co-chaperones remove sHsps from the assembly. This process might explain why HSPA1A/B is more abundant in UPODs than HSPB1. Overall, our results suggest that UPODs may be generally targeted by sHsps in collaboration with Hsp70 to modulate the formation of UPODs and refold IFPs. This process does not seem to be restricted to barnase UPODs, as HSPB1 has been shown to suppress mutant SOD1 aggregation *in vitro*, increase survival when overexpressed with Hsp70 in an ALS cell model, and inhibit early stages of disease progression in ALS mouse models (Patel et al., 2005; Sharp et al., 2008; Yerbury et al., 2013).

The shared chaperone regulation pathway between the model protein barnase and a human disease related IFP suggests that features that influence recruitment into UPODs formed by barnase variants are also likely to be transferrable to other IFPs. Of interest is the observation that proteins can be recruited to UPODs based on a shared grammar for interactions of cellular proteins with the unfolded states of the phase separating IFP. A similar result was found by Wear et al. They showed that Httex1 with expanded polyglutamine tracts recruit proteins with long IDRs into its deposits (Wear et al., 2015). Deletion of the long IDRs in two of the recruited proteins decreases colocalization with Httex1. These results suggest that IDR-IDR interactions between Httex1 and other cellular proteins may lead to sequestration and subsequent loss-of-function of the recruited proteins.

Our results suggest that sequestration based on shared sets of interactions will be specific to the phase separating IFP. Specifically, the distribution and type of stickers in the unfolded states of an IFP can contribute to distinct types of heterotypic interactions. This result suggests that the identification of stickers within the unfolded states of an IFP will be important if we are to modulate aberrant homotypic interactions that drive UPOD formation and heterotypic interactions that lead to sequestration and potential loss-of-function of the recruited proteins.

Additionally, the recruitment of proteins based on shared interaction grammars imply that the relevant residues must be accessible for heterotypic interactions. Residues may be accessible if they are part of an IDR. However, 66 of the top 94 differently enriched proteins (70%) do not contain an IDR of length greater than 50. Instead, residues may be accessible if they are sequestered in UPODs before they have the chance to fold. Olzcha et al., showed that newly synthesized proteins preferentially interact with a synthetic beta-sheet protein that forms deposits via homotypic interactions (Olzscha et al., 2011). If newly synthesized proteins are also preferentially recruited to UPODs, then it may imply that proteins that require long time scales to fold or the help of many chaperones will be particularly susceptible to recruitment into aberrant UPODs. Interestingly, both Hsp70 and TRiC can work together for co-translational folding of substrates (Stein et al., 2019). Thus, the recruitment of these chaperones to UPODs may further increase the population of unfolded or improperly folded newly synthesized proteins.

Kaganovich et al., previously identified two types of protein quality control compartments they named the insoluble protein deposit (IPOD) and juxtanuclear quality control compartment (JUNQ) (Kaganovich et al., 2008). Interestingly, polyglutamine containing proteins were shown to form IPOD structures (Kaganovich et al., 2008), whereas other misfolded proteins, such as SOD1 destabilizing variants, formed JUNQs (Polling et al., 2014). Reversibly misfolded proteins were proposed to be delivered to the JUNQ compartment for refolding, whereas terminally misfolded proteins go into the IPOD compartment for sequestration. The colocalization of the I25A, I96G barnase variant with SOD1 and the enrichment of HSPA1A/B in barnase UPODs suggests that UPODs may be equivalent to JUNQ compartments (Weisberg et al., 2012). However, further compositional profiling and staining for JUNQ markers will be necessary to determine whether UPODs and JUNQs are equivalent or distinct. If the two compartments are equivalent, then our compositional profiling experiments would suggest the compositions of JUNQs are unique to the sticker grammars of unfolded / misfolded IFP(s) that drive its formation. Differences in composition could lead to differences in cell-specific stresses. Therefore, determining the relationship between UPODs and JUNQs is important for understanding how the cell manages unfolded / misfolded IFPs and the progression of disease.

## ACKNOWLEDGMENTS

We are grateful to our colleagues and collaborators Chloe Gerak, Mina Farag, Beatriz Ferreira-Gomes, Marta Frigole-Vivas, Paul Gooley, Matthew King, Tanja Mittag, Ammon Posey, and Min Kyung Shinn for helpful discussions. The contributions of Y H.C., A.R.O., D.C., and D.M.H., were supported by grants APP1161803 and APP1154352 from the National Health and Medical Research Council of Australia, and DP170103093 from the Australian Research Council awarded to D.M.H. The contributions of K.M.R. and R.V.P were supported by grants from the US National Institutes of Health (5R01NS056114) and the US Air Force Office of Scientific Research (FA9550-20-1-0241). S.E.R. is a Royal Society Research Professor (RSRP\R1\211057) and is funded by the Wellcome Trust (204963).

## AUTHOR CONTRIBUTIONS

D.M.H. and R.V.P conceived of the project. Y H.C, D.C., and A.R.O. performed in the *in vitro* and in cell measurements. K.M.R., R.V. P., and D.M.H., designed the analysis. K.M.R. performed the simulations and computational analysis. Y.M. and D.A.B. helped generate estimates of stabilities for the barnase variants. S.E.R. and D.C. helped with critical thinking and variant design. K.M.R. designed the figures, and K.M.R., Y H.C, D.C., and A.R.O. made the figures. K.M.R., D.M.H., and R.V.P. wrote and edited the first full drafts of the manuscript. K.M.R., Y H.C., D.C., A.R.O., R.V.P., and D.M.H., collaborated extensively on the final rounds of editing and all authors were involved in revising the manuscript. R.V.P. and D.M.H. secured funding.

## STAR Methods

### Cell culture and imaging

Mouse Neuro2a and human HEK293T cells were used in this study. Neuro2a and HEK293T cells were maintained in opti-MEM and Dulbecco’s Modified Eagle Medium (DMEM) respectively, supplemented with 10% v/v foetal bovine serum and 2 mM L-glutamine (Thermo Fisher Scientific) in a humidified incubator at 37 °C and 5% v/v atmospheric CO_2_. For all imaging experiments, cells were plated at 3×10^4^ cells per well in 8-well µ-slides (Ibidi) and transfected using Lipofectamine 3000 (Thermo Fisher Scientific) as per the manufacturer’s protocol. In the case of HEK293T cells, plates were pre-coated with poly-L-lysine to aid adhesion. Imaging was conducted on a Leica TCS SP5 Confocal microscope using a HCX APO CS 63 × 1.40 Oil objective lens unless stated otherwise.

For optoDroplet experiments, cells were stained 24 h post-transfection, with Hoechst 33342 at 20 µM for 20 min at 37 °C, washed and imaged in Hank’s Balanced Salt Solution (HBSS). mCherry fluorescence was imaged (561 nm excitation, 600-650 nm emission) prior to optoDroplet activation, followed by photoactivation with the 488 nm laser for 60 s at a laser intensity of 30%. mCherry and Hoechst fluorescence (excitation 405 nm, emission 420-540 nm) were then imaged immediately after activation. Droplet disassembly was observed post-activation by time-lapse imaging of mCherry fluorescence every 60 s for 15 min.

For VER-155008 treatment with optoDroplet expression experiments, Neuro2A cells were transiently transfected with barnase L14A in the optoDroplet construct for 24 h. After transfection, transfection media was removed and cells were incubated with opti-MEM containing 0, 1, 5, 10 µM VER-155008 dissolved in dimethyl sulfoxide (DMSO) for 2 or 4 h. After treatment, drug-treatment media was removed, and cells were washed twice with PBS before being stained with Hoechst 33342 and imaged on the confocal in Hank’s Balanced Salt Solution (HBSS) and imaged on the confocal. Imaging for VER-155008-treated cells were conducted on a Zeiss LSM900 confocal microscope using a Plan-Apochromat 40 × 1.2 oil objective lens. With the exception of optoDroplet activation at 20% laser intensity, imaging parameters were kept the same as described above.

For chaperone optoDroplet experiments, Neuro2A cells coexpressed either opto-barnase with HSPA1A and DNAJB1, opto-barnase with HSPA1A or DNAJB1 and emerald (Y66L) or opto-barnase with emerald (Y66L). The cells expressing three constructs were transfected at a concentration ratio of 1:1:1 and cells expressing the opto-barnase with emerald (Y66L) were transfected at a 1:2 ratio. Emerald (Y66L) was used as an inert control protein to ensure the same amount of opto-barnase DNA was being added to the cells while maintaining the recommended DNA amount for lipofectamine transfection. Imaging was carried out as described above for optoDroplet experiments.

For immunofluorescence, cells were fixed 24 h post-transfection in 4% w/v paraformaldehyde for 15 min at room temperature. Cells were then permeabilized with 0.5% v/v Triton X-100 in phosphate buffered saline (PBS) for 20 mins at room temperature. Samples were blocked in 5% w/v bovine serum albumin in PBS for 1 hour at room temperature followed by staining with anti-V5 antibody (1:250 dilution, Abcam cat# ab27671) or anti-HSPA1A (1:100 dilution, Abcam cat#ab5439) diluted in PBS containing 1% w/v bovine serum albumin and 0.3% v/v Triton X-100 overnight at 4°C. Samples were then incubated in goat anti-mouse Cyanine5 (1:500) (Life technologies cat# A10524) diluted in PBS for 30 mins at room temperature. Finally, cell nuclei were stained with Hoechst 33342 at 20 µM for 20 min at 37 °C. Cyanine5 fluorescence was imaged using 633 nm excitation and 695-765 nm emission and Hoechst using 405 nm excitation and 410-450 nm emission.

### Constructs

The sequence for the DDX4-mCherry-Cry2 optoDroplet construct, which was based on the work of Shin et al. (Shin et al., 2017), was synthesised (Thermo Fisher Scientific) and cloned into the pTriEx4 expression vector by restriction cloning using BamHI and XhoI restriction enzymes. Barnase optoDroplet constructs were generated by PCR amplification, restriction digestion using BamHI and SacI restriction enzymes, and ligation to replace the DDX4 with barnase variants. Barnase sticker variants were synthesised (Genscript) in the pTriEx4 optoDroplet expression vector. Additional barnase variants were synthesized as cassettes (GenScript) and cloned into the pTriEx4 optoDroplet expression vector using BamHI and SacI restriction enzymes. Barnase and SOD1 were cloned into the pTriEx4 FRET vectors using the FastCloning strategy (Li et al., 2011) where the inserts and vector were PCR amplified with overlapping primers, template plasmids were digested with the methylation-sensitive restriction enzyme DpnI, and the product was directly transformed in chemically competent DH5α cells. Hsp40 and Hsp70 constructs were prepared as described previously (Ormsby et al., 2013). V5-tagged chaperone proteins were overexpressed from pcDNA5/FRT/TO V5 DNAJB1 and pcDNA5/FRT/TO V5 HSPA1A provided as gifts from Harm Kampinga (Hageman and Kampinga, 2009) via Addgene. Httex1-72Q fused to mCherry in the pGW1 vector were prepared as previously described (Arrasate et al., 2004) and kindly provided by Steven Finkbeiner. All constructs were verified by sequencing.

### Image Analysis

Representative confocal micrographs including cell outlines were manually produced using FiJi (Schindelin et al., 2012). The brightness and contrast of individual images were adjusted to maximise the visible range of fluorescence intensity across constructs with different ranges of expression. Additional quantitative analyses on unmodified images were carried out using custom scripts written in the python programming language. Cells and nuclei were first automatically segmented using the Cellpose package (Stringer et al., 2021), and segmentation was manually inspected for quality control using Napari (Tyson et al., 2021). Cells on the image boundary, those that did not contain a nucleus, or those that were associated with more than one nucleus, were removed from subsequent analyses. Pixel coordinates were then extracted for the individual whole-cell and nuclei segmentation masks. Coordinates of nuclei were excluded from whole-cell coordinates to yield cytoplasmic pixels. For immunofluorescence experiments, compartment fluorescence was calculated as the mean intensity of pixels in the nucleus or cytoplasm respectively. Pixel intensities for individual cells were saved as csv files for further analysis as indicated below.

### Extraction of *c*_sat_ values for optoDroplet formation of barnase

Using raw pixel data extracted from the confocal fluorescence micrographs, pixel intensities of all cells were first converted to natural log space. Cells in which greater than 25% of the pixels have the max intensity were then removed. For the remaining cells, pixel intensity histograms were generated using the data obtained prior to activation to identify the dominant peak. Since cells should have approximately uniform intensity before activation, the histograms were fit to a Gaussian distribution to filter out pixels whose intensities were not numerically similar to the mean intensity. Specifically, the Gaussian fit was used to identify the maximum frequency of the histogram and the mean intensity. The width of the distribution was then determined by finding the first instances of 20% of the maximum frequency on either side of the mean intensity. All pixels that did not fall within the intensity bins bound by this filter were removed. We also removed all pixels that were already at the maximum intensity before activation. This filtering process accounts for the fact that before activation cells should have relatively uniform intensities. The positions of the filtered pixels were then used to extract the relevant pixels from the data obtained after activation. Raw intensity histograms of the before and after activation data were then created using the filtered pixels. We further removed cells in which histograms had data in less than or equal to five bins and had fewer than 100-pixel positions. These filters make sure that there is enough data for a Gaussian fit of the histograms to be reasonable. Then, the before activation histogram was fit to a single Gaussian and the mean intensity before activation (*I*_dil,before_) was collected. This mean intensity should be proportional to the total concentration of barnase. The after-activation histogram was then fit to a model that is a mixture of two Gaussians. The premise is that if phase separation occurs there should be low intensity and high intensity peak, where the mean of the low intensity peak (*I*_dil,after_) is proportional to the concentration of barnase in the dilute phase and the mean of the high intensity peak (*I*_den,after_) is proportional to the concentration of barnase in the dense phase. We restrict the fit such that the *I*_dil,after_ ≤ *I*_dil,before_, since the concentration of barnase in the dilute phase should not be greater than the total barnase concentration. Any cells in which *I*_dil,after_ or *I*_dil,before_ were less than zero or the *R*^2^ value for the after-activation fit was less than 0.85 were removed. For the remaining cells, *I*_dil,after_ was then collected.

Next, if there is no phase separation *I*_dil,before_ and *I*_dil,after_ should follow a one-to-one correspondence, i.e., *I*_dil,before_ ≈ *I*_dil,after_. It follows that the *c*_sat_ for phase separation should correspond to the intensity at which this one-to-one correspondence no longer holds. To extract the *c*_sat_, we first removed outliers in the *I*_dil,before_ versus *I*_dil,after_ plots using three steps. Before any step was performed two threshold values were set. The value *x*_1to1_ = 1000 corresponds to *I*_dil,before_ -*I*_dil,after_ threshold for a cell to still be considered within the one-to-one regime. Then, for all cells outside of this regime, *m*=0.9*mean(*I*_dil,before_ - *I*_dil,after_) was calculated. Here, *m* is used to distinguish between cells slightly outside the one-to-one regime or largely outside this regime. The first step to remove outliers consisted of removing cells that fell off the diagonal even though other cells in this intensity regime showed one-to-one behaviour. Next additional outliers were removed using Cook’s distance (Cook, 1977). Specifically, any cells that corresponded to *I*_dil,before_ - *I*_dil,after_ < *m* were fit using a linear regression model and any cells that were 5*mean(Cook’s distance for all points) were filtered out. Finally, cells well outside the one-to-one regime (*I*_dil,before_ - *I*_dil,after_ ≥ *m*) were fit using a linear regression model and any cells that were 5*mean(Cook’s distance for all points) were filtered out.

Once outliers were removed, each barnase variant was checked for whether they had at least three off diagonal cells (*I*_dil,before_ - *I*_dil,after_ ≥ *m*) so a fit for *c*_sat_ could be performed. Barnase variants that did not satisfy this cut-off were taken as not undergoing phase separation. For the remaining barnase variants, 50 bootstrapping trials were performed with the sample number corresponding to 0.9 times the number of cells corresponding to *I*_dil,before_ - *I*_dil,after_ ≥ *m*. Then these cells and the cells corresponding to the one-to-one regime were systematically split into two data sets to extract the optimal fit of *c*_sat_. Specifically, all cells corresponding to *I*_dil,before_ ≥ *splitVal* were fit using a linear regression that crossed the one-to-one line at *splitVal*. The *splitVal* that minimized both the sum of squares due to error and 1-*R*^2^ was taken to be the *c*_sat_ for phase separation.

### Fitting of *c*_sat_ versus 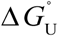

The fraction of unfolded proteins for a given 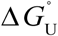 of unfolding is given by:

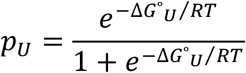

where 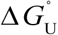 is the standard state free energy of unfolding, *R* is the gas constant (8.131 J/mol-K) and *T* is the temperature (293 K). However, even variants with 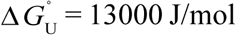, which equates to >99% folded molecules, can undergo phase separation. This suggests barnase variants may be more unstable in cells than their *in vitro* 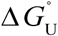 values imply. Thus, we define the fraction of unfolded proteins for the shifted 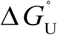 as follows:

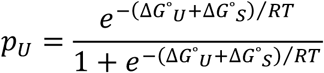

where Δ*G°*_S_ denotes the constant offset. We then fit the extracted *c*_sat_ values assuming phase separation occurs at a critical unfolded concentration, using *c*\* = *c*_sat_×*p*_U_.

### *c*\* confidence interval

To identify hydrophobic and hydrophilic blobs in the barnase sequences we utilized the method of Lohia et al. (Lohia et al., 2019). Briefly, the average scaled Kyte-Doolittle hydrophobicity score (0 to 1) was calculated over three residue windows. Four or more contiguous windows with a score > 0.37 is considered a hydrophobic blob, whereas four or more contiguous windows with a score of ≤ 0.37 is considered a hydrophilic blob. We define the change in blobs from wild type as the sum of the magnitude of the decrease in size of hydrophobic blobs and the increase in size of hydrophilic blobs. To determine the confidence interval for *c*\* a picking weight for each barnase variant sequence was determined by (10-(|decrease in size of hydrophobic blobs| + increase in size of hydrophilic blobs))/10, to ensure that sequences that had limited change in blobs were picked more often. Then 1000 bootstrapping trials were performed selecting 10 sequences each time based on these weights. Each trial was fit as described above to extract Δ*G°*_S_ and *c*\*. Then, the mean and standard deviations of these values were calculated. The interval corresponds to plotting *c*_sat_ = *c*\*/ *p*_U_ with (mean(*c*\*)+std(*c*\*), mean(Δ*G°*_S_)-std(Δ*G°*_S_)) and (mean(*c*\*)-std(*c*\*), mean(Δ*G°*_S_)+std(Δ*G°*_S_)).

### Computational mutagenesis study predicting Δ*G°*_U_

To assess the effect of mutations on the stability of barnase, the x-ray structure of barnase (PDB ID: 1A2P, resolution of 1.5 Å) was obtained from Protein Data Bank. We further processed the 3D structure by removing redundant chains, ions, water molecules and alternative conformations of residues 28, 31, 38, 85 and 96. The stability changes upon mutations, measured as the change in Gibbs Free Energy (ΔΔG in kcal/mol), were predicted using “Calculate Mutation Energy (Stability)” in Discovery Studio 2018 (https://www.3ds.com/products-services/biovia/products/molecular-modeling-simulation/biovia-discovery-studio/) with preliminary minimization of wild-type structure. The results of single and double mutations were used to build a transformation model to adjust the predicted ΔΔG of high multiple mutations (up to 8 mutations per case).

### Atomistic simulations

Atomistic simulations were performed using the ABSINTH implicit solvation model and forcefield paradigm (Vitalis and Pappu, 2009) as implemented in the CAMPARI simulation engine (http://campari.sourceforge.net). Simulations were performed using a parameter set based on abs3.2_opls.prm(https://github.com/kierstenruff/RUFF_CHOI_unfolded_protein_phase_separation). Each simulation was performed in a spherical droplet of radius 150 Å (barnase, SOD1) or 200 Å (HSPB1, GSTP1, PRDX1, RPS28, H3-3A) at 335 K. The droplet radius was increased for the additional IFPs given that the sequence length of many of these IFPs is ∼200. Additionally, counterions and an excess of 5 mM NaCl were modelled explicitly. Each Metropolis Monte Carlo simulation comprised 10^7^ equilibration steps and 5.15 × 10^7^ production steps. For each construct, we performed five independent simulations. To model the unfolded state, simulations were started from completely random structures. For Httex1-49Q, we reanalyzed simulations performed at 335 K from the work of Warner et al., (Warner et al., 2017). Reference Flory Random Coil (FRC) simulations were performed as described in Holehouse et al., (Holehouse et al., 2015). Briefly, backbone and side-chain dihedral angles are randomly drawn from previously generated dipeptide simulations to construct ensembles in which chain-chain and chain-solvent interactions are counterbalanced.

### Identifying stickers from atomistic simulations of unfolded states

The mean contact probability for each residue was calculated using the SOURSOP analysis package https://github.com/holehouse-lab/soursop. Here, the probability that a residue is in contact with another residue is averaged over all residues, excluding the nearest and second nearest neighbor contacts. The cut-off for a contact was set to 5 Å. Given that the mean contact probability will be dependent on both sequence length and amino acid sequence, we also calculated the mean contact probability for each construct from the reference FRC simulations. Strong stickers should prefer chain-chain interactions and thus have a larger mean contact probability than what is observed in the corresponding FRC simulation in which chain-chain and chain-solvent interactions are counterbalanced. Therefore, we define a strong sticker by a mean contact probability greater than the maximum mean contact probability from the corresponding FRC simulation.

To identify the type(s) of residues are most likely stickers we grouped residues into six categories. The aliphatic residues include Ala, Ile, Leu, Met, and Val; the aromatic residues include Phe, Trp, and Tyr; the unique residues include Cys and Pro; the acidic residues include Asp and Glu; the basic residues include His, Lys, and Arg; finally, the polar residues include Gly, Asn, Gln, Ser, and Thr. We calculated the fraction of total mean contact probability for each type and compared it to the expected total mean contact probability based on the number of residues in the sequence of that given type. Residue types featuring a high fraction of mean contact probability that is also greater than what we expect based on their numbers of occurrence within the sequence are likely going to be the predominant stickers.

### Flow cytometry

HEK293T cells were plated on a poly-L-lysine-coated 24-well plate (Falcon) at a density of 7.5 × 10^4^ cells per well and transfected with lipofectamine 3000 as per manufacturer’s protocol. Following 48 h after transfection, HEK293T cells were washed once with PBS and detached by gentle pipetting and transferred into a U-bottom microplate. Flow cytometry was performed as described previously (Wood et al., 2018). Flow cytometry data were processed with FlowJo (Tree Star Inc.) to exclude un-transfected cells and cell debris and compensate the Venus channel to remove bleed-through from the mTFP1 and FRET channels. The mTFP1, Venus and FRET data were exported as csv files for further analysis. Barnase A_50_ were calculated as previously described (Wood et al., 2018).

### LC-MS/MS sample preparation and analysis

#### Sample preparation for proteomics

Four biological replicates were used for each sample group. 1.7×10^6^ HEK293T cells were seeded in 25 cm^2^ flasks 24 h before transfection. Cells were transfected with barnase variants in the FRET construct. Cells were transiently transfected as per manufacturer’s protocol, the transfection media was removed 6 h post-transfection and cells were incubated for a total of 48 h, including the 6 h incubation in the transfection media. Post-transfection, cells were washed and harvested in PBS by gentle pipetting and incubated in 200 µg/ml digitonin dissolved in PBS for 20 min at room temperature to remove diffuse cytosolic proteins. Cells were again pelleted, the supernatant was collected as the soluble fraction and the pellet was resuspended with RIPA lysis buffer (150 mM NaCl, 50 mM Tris-HCl, pH 8.0, 1% NP-40, 0.5% sodium deoxycholate, 0.1% SDS, Complete EDTA-free protease inhibitor (Roche), 25 U/ml benzonase) and incubated for 10 min at room temperature. Cells were further lysed mechanically by vortexing and pipetting. 8 M urea dissolved in 50 mM Tris-HCl (pH 8.0) was added to the lysates to a final concentration of 4 M and incubated at room temperature for 15 min, then sonicated for 15 min to solubilize the aggregates. The lysates were centrifuged (21000 ×g; 15 min; 4 °C) to remove cell debris and any aggregated proteins that were not solubilized and the supernatant containing the solubilized aggregates were collected. The protein concentration for each sample was determined using bicinchoninic acid assay (BCA), as per manufacturer’s protocol (Thermofisher) and 100 μg of each sample was incubated in ice-cold acetone overnight at –20°C. The acetone-precipitated samples were pelleted (20000 ×g; 30 min; 4°C), acetone was removed, and pellets were dried, ensuring all the acetone had evaporated. The protein pellets were resuspended and incubated in 50 mM TEAB, 8 M urea (pH 8.0) for 30 min at 37°C. Tris(2-carboxyethyl)phosphine (TCEP) was added to a final concentration of 10 mM and samples were incubated for 45 min at 37°C to reduce the proteins, followed by the addition of iodoacetamide to a concentration of 55 mM and incubated for a further 45 min at 37°C to alkylate the proteins. The samples were diluted to a final concentration of 1 M urea and digested overnight at 37°C with 2.5 µg of trypsin. The trypsin-digested peptides were acidified with pure formic acid to a final concentration of 1% (v/v) and a sample cleanup using a solid phase extraction (SPE) method was performed. SPE cartridges (Waters/Oasis) were first equilibrated with 80% acetonitrile, 0.1% trifluoroacetic acid (TFA) followed by 0.1% TFA prior to loading samples onto the column. Bound peptides were washed twice with 1.5 ml 0.1% TFA and then eluted in 800 µl 80% acetonitrile, 0.1% TFA. Peptide samples were vacuum dried using a SpeedVac vacuum concentrator and resuspended in double-distilled water for tandem mass tag (TMT) labelling.

#### TMT labelling for proteomics

TMT labelling was conducted based on the manufacturer’s protocol (Thermofisher). 1 M TEAB and acetonitrile was added to each sample to a concentration of 30% acetonitrile and TMT labelling reagents were resuspended in acetonitrile. TMT label reagents were added to samples in an 8:1(m/m) ratio and incubated at room temperature for 1 h. Samples were mixed by vortex at regular intervals during incubation. The reaction was quenched with 8 µl of 5% hydroxylamine for 15 min at room temperature. Each label was combined in a 1:1 mass ratio for MS analysis.

#### Mass spectrometry data acquisition and analysis

10 µg of TMT-labelled peptide mixtures were lypophilised using a SpeedVac vacuum concentrator and resuspended to a final concentration of 0.5 µg/µl in 2% (v/v) acetonitrile, 0.05% (v/v) TFA. Peptides were analysed by nanoESI-LC-MS/MS using the Thermo Orbitrap Q Exactive Plus mass spectrometer (Thermofisher) equipped with a nanoflow reversed-phase-HPLC (Ultimate 3000 RSLC, Dionex) fitted with an Acclaim Pepmap nano-trap column (Dionex—C18, 100 Å, 75 μm× 2 cm) and an Acclaim Pepmap RSLC analytical column (Dionex—C18, 100 Å, 75 μm× 50 cm) by the University of Melbourne Mass Spectrometry and Proteomics facility. 0.6 µg of the TMT-labelled peptide mixture was loaded onto the enrichment (trap) column at an isocratic flow of 5 µl/min of 2% acetonitrile containing 0.1% (v/v) formic acid for 5 min. The enrichment column was then switched in-line with the analytical column. The eluents used for the liquid chromatography were 0.1% (v/v) formic acid, 5% (v/v) DMSO for solvent A and 0.1% formic acid (v/v), 5% (v/v) DMSO in acetonitrile for solvent B, flowed at 300 nl/min using a gradient of 3–22% solvent B in 90 min, 22–40% solvent B in 10 min and 40–80% solvent B in 5 min then maintained for 5 min before reequilibration for 8 min at 3% B prior to the next analysis. All spectra were acquired in positive ionization mode with full scan MS acquired from *m/z* 300-1600 in the FT mode at a mass resolving power of 120 000, after accumulating to an AGC target value of 3.0e^6^, with a maximum accumulation time of 25 ms. The RunStart EASY-IC lock internal lockmass was used. Data-dependent HCD MS/MS of charge states > 1 was performed using a 3 s scan method, isolation width of 0.7 m/z, at a normalised AGC target of 200%, automatic injection time, a normalised collision energy of 30% and with spectra acquired at a resolving power of 30000 (TurboTMT activated). Dynamic exclusion was used for 20 s.

Data analysis was conducted using MaxQuant (version 2.0.1.0.) and database searches were conducted using the Swissprot Homo sapiens database (accessed on 6^th^ July 2021, 20371 entries) with the additional barnase WT, 8×A, 9S, mTFP1 and Venus proteins. The search was conducted with 20 ppm MS tolerance, 0.5 Da MS/MS tolerance and 2 missed cleavages allowed. Oxidation (M) and acetyl (Protein N-term) variable modifications were allowed and a fixed modification for carbamidomethyl (C) was used for all samples. The false discovery rate was set at 1% for both peptides and proteins. Peptide and protein abundances were normalized by the total abundance of all identified proteins in each sample group. Proteins that were identified in less than or equal to three of the replicates in any of the sample groups were excluded from analysis. For proteins with missing values in one out of the four replicates, the missing value was filled with the mean abundance of the protein in the sample group. The initial data cleanup and normalization was conducted in python and the source code can be accessed via https://github.com/kierstenruff/RUFF_CHOI_unfolded_protein_phase_separation.

Multivariate analysis of proteomics data was conducted using the online software Metaboanalyst (Pang *et al*., 2021). A one-way ANOVA with a p-value cutoff of 0.05 using a Fisher’s LSD posthoc test was conducted to determine the proteins that were significantly differently enriched in the UPODs of the barnase variants. Gene ontology search was conducted using PANTHER database (Mi *et al*., 2021). For visualization of the abundance z-scores a smoothing procedure was performed in which the mean value was scaled based on its p-value following a one-sample t-test (Cox et al., 2022).

#### Assignment of functional categories

Functional categories in **Figure S5E** were determined by mining the Gene ontology (biological process) and Gene ontology (molecular function) categories using UniProt, as well the function listed in the Human Protein Atlas for each of the top 94 differentially enriched proteins (**Table S5**)(Consortium, 2020; Uhlén et al., 2015).

#### Sequence Feature Analysis

Ninety-one sequence features were examined for each of the top 94 differently enriched endogenous proteins by a one-way ANOVA in the proteomics dataset. Many of the sequence features are the same as those identified by Zarin et al., to be important for the molecular function of disordered regions (Zarin et al., 2019). We added additional sequence features that have been shown to be important for function or phase behavior of disordered regions. We focus on these features as they are likely to be important for interactions in the unfolded state as well. The sequence features can be split into 2 distinct categories: (1) patterning and (2) composition. For the patterning features a modified version of NARDINI was deployed (Cohan et al., 2022). The specific modifications were as follows: we generated 10^3^ scrambled sequences per variant rather than the 10^5^ and each distribution is not fit to a gamma distribution, as these changes did not have large effects on the overall outcome. Additionally, we set *g*, the number of residues in a sliding window, to be 5 and 6 and take the mean of these results. In NARDINI, the residue groups are defined as follows: pol ≡ (S, T, N, Q, C, H), hyd ≡ (I, L, M, V), pos ≡ (K, R), neg ≡ (E, D), aro ≡ (F, W, Y), ala ≡ (A), pro ≡ (P), and gly ≡ (G). Z-scores below zero imply that the original sequence is more well-mixed with respect to the residue groups compared to the scrambled sequences. Z-scores above zero imply that the original sequence is blockier with respect to the residue groups compared to the scrambled sequences.

The composition features consist of 55 features including: the fraction of each amino acid (20); the fraction of positive, negative, polar, aliphatic, aromatic, charged, chain expanding, disorder promoting, and (R, Y) residues (9 in all); the ratio of Rs to Ks and Es to Ds (2); the net charge per residue, the mean hydrophobicity, the isoelectric point, the polyproline-II propensity, and the number of (R, Y) residues (5). Additionally, we included patch features which are calculated as the fraction of the sequence that is made up of all patches of a particular amino acid or RG. Here, a patch must have at least four occurrences of the residue or two occurrences of RG and cannot extend past two interruptions. This leads to an additional 19 features given that there were no M or W patches in the proteomics dataset. localCIDER was used to extract a majority of the composition sequence features (Holehouse et al., 2017). To calculate z-scores, the composition features were calculated over all 2269 mapped proteins in the proteomics dataset. Then the z-score for each of the top 94 proteins was calculated using the mean and standard deviation of 2269 proteins.

Finally, we also added an abundance feature to our analysis. Here, the mean and standard deviation of abundance in UPODs was calculated over all barnase variants and replicates for all 2269 mapped proteins in the proteomics dataset. These values were used to calculate the abundance z-score for each of the top 94 proteins. Together, this yielded 92 distinct z-scores for each of the top 94 proteins.

We extracted the set of proteins that are significantly enriched in a specific set of barnase UPODs compared to either all remaining barnase UPODs (Rest) or a different set of barnase variants using a *post hoc* Fisher’s least significant difference (LSD) test following an ANOVA test. To determine which features were distinct to a set of proteins enriched in a specific barnase UPOD(s), we compared the z-score distribution of the 92 features in each enriched set to the z-score distribution of the remaining top 94 proteins using the two-sample Kolmogorov-Smirnov test. If the p-value was less than 0.05 the signed log(p-value) was recorded to identify the significant features of recruited proteins associated with a specific UPOD(s). Here, the log(p-value) was positive if the median of the z-score distribution associated with the set of proteins that are significantly enriched in a specific barnase UPOD(s) was greater than the median of the z-score distribution for the remaining top 94 proteins.

### Disorder Analysis

We used a previously generated in house disorder database. The database was generated using the Swissprot Homo sapiens database (accessed in May 2015, 20882 entries) (Consortium, 2020). The predicted disorder for each sequence was determined by running MobiDB (Piovesan et al., 2020). A residue is considered disordered if the consensus prediction labeled it as disordered.

## Data and code availability

The original data collected and scripts used for all analyses can be downloaded from GitHub (https://github.com/kierstenruff/RUFF_CHOI_unfolded_protein_phase_separation). The RAW files have been deposited via the <u>PRIDE</u> partner repository to the <u>ProteomeXchange Consortium</u> (Perez-Riverol et al., 2018) with the dataset identifier PXD033716. Additionally, select summary data are provided in Tables S1-5.

