## Supplemental Material for "Sequence grammar underlying unfolding and phase separation of globular proteins"

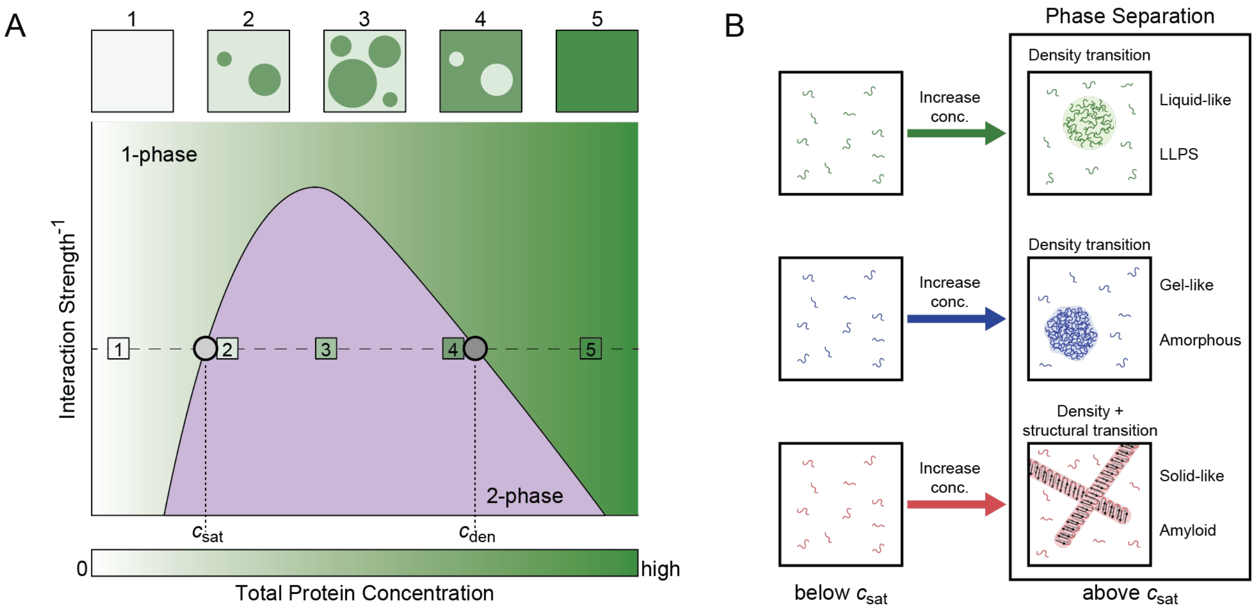


**Figure S1. Phase separation is a density transition that occurs above the saturation concentration, $c_{sat}$. Related to Figure 1.** (A) Schematic of homotypic phase separation. For a given (interaction strength)$^{-1}$ (dashed line), the system contains only one phase when the total protein concentration is less than $c_{sat}$ (box 1). At a total protein concentration equal to $c_{sat}$, the system separates into a protein-deficient phase (dilute phase) and a protein-rich phase (dense phase). Above $c_{sat}$ and below $c_{den}$, the concentration of the protein in the dilute phase is given by $c_{sat}$ and the concentration of the protein in the dense phase is given by $c_{den}$. The volume of the dense phase increases as the total concentration of protein increases (see boxes 2-4). Finally, above $c_{den}$ the system returns to a single phase. (B) The schematic shows how crossing $c_{sat}$ gives rise to different types of coexisting phases, distinguished by the material properties of the dense phase.

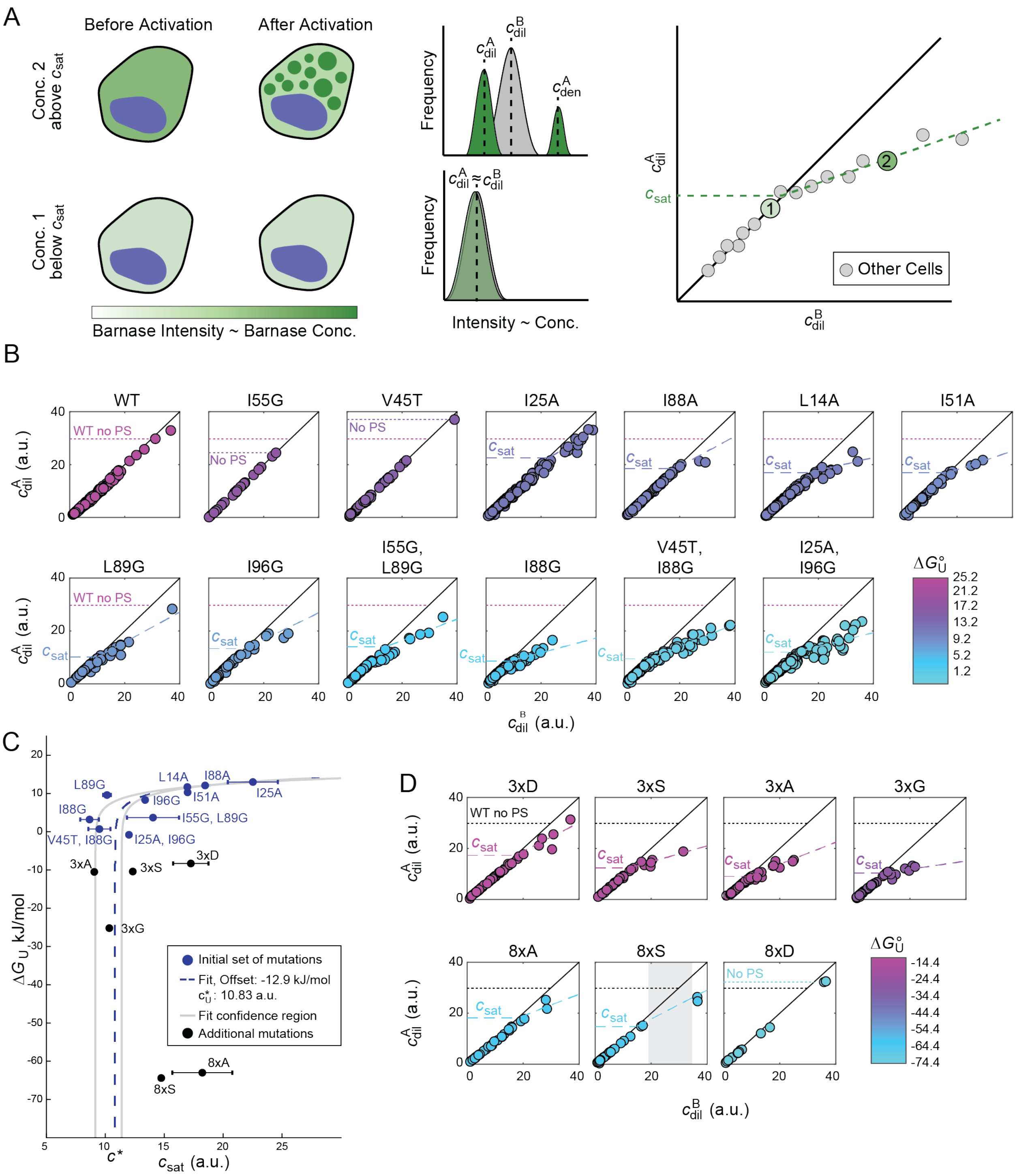


**Figure S2. Extraction of $c_{sat}$ for barnase variants. Related to Figure 3.** (A) Schematic depicting how the fluorescence intensities before and after light activation can be used to extract the protein concentration (in arbitrary units) of the dilute phase before activation, $c_{dil}^B$, and the protein concentration of the dilute phase after activation, $c_{dil}^A$. Darker green colors imply higher concentrations of protein. For cells that do not undergo phase separation upon light activation, $c_{dil}^B \approx c_{dil}^A$. For cells that undergo phase separation, $c_{dil}^A$ will be less than $c_{dil}^B$. Thus, the $c_{sat}$ corresponds to where $c_{dil}^A$ no longer equals $c_{dil}^B$. Blue blobs indicate the nucleus in cells. (B) $c_{dil}^B$ versus $c_{dil}^A$, in arbitrary units, for the initial barnase variants examined in Figure 3. Here, each circle is a cell. Dotted lines denote that the $c_{sat}$ must be larger than this value as no phase separation was observed up to this concentration. Dashed lines denote the fitted $c_{sat}$ (see **STAR Methods**). (C) $c_{sat}$ for optoDroplet formation, in arbitrary units (a.u.), versus $\Delta G^\circ_U$. Error bars indicate standard deviations across 50 bootstrapped trials. Dashed line corresponds to a fit of the initial barnase construct data (blue circles) to a model that assumes only crossing a critical concentration of unfolded proteins is necessary for phase separation. Fit assumes that barnase is more unfolded in the cell than predicted by $\Delta G^{\circ}_{U}$ = -12.9 kJ/mol with $c^*$ = 10.83 a.u. Gray lines denote the fitted confidence interval determined by bootstrapped trials of the variants, where the variants were picked based on the degree to which they modulated the hydrophobic and hydrophilic blobs from WT (**STAR Methods**). Black points show results for additional, more extreme barnase variants in which effectively all molecules should be in the unfolded state. (D) $c_{dil}^{B}$ versus $c_{dil}^{A}$, in arbitrary units, for the extreme barnase variants examined in Figure 3. Grey box in 8xS shows a void in concentrations examined which makes the exact $c_{sat}$ value for this variant unclear.

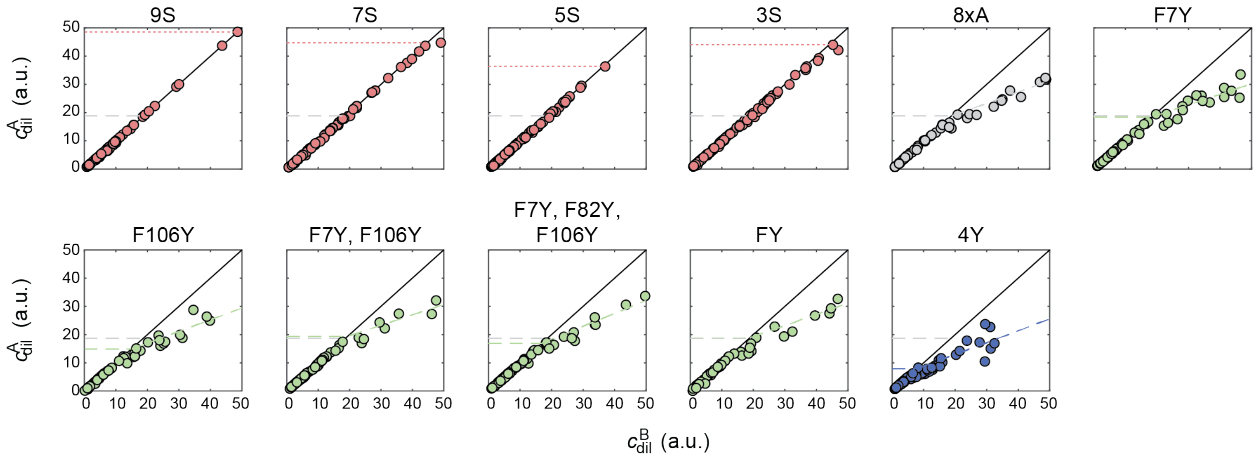


**Figure S3. Extraction of $c_{sat}$ for barnase sticker variants. Related to Figure 4.** Dilute phase concentration before, $c_{dil}^{B}$, versus after activation, $c_{dil}^{A}$, in arbitrary units. Here, each circle is a cell. Dotted lines denote the $c_{sat}$ must be larger than this value as no phase separation was observed up to this concentration. Dashed lines denote the fitted $c_{sat}$ (see **STAR Methods**). Red points correspond to variants that reduce the number of stickers, grey to the template variant, green to variants that replace F with Y, and blue to a variant that increases the number of stickers.

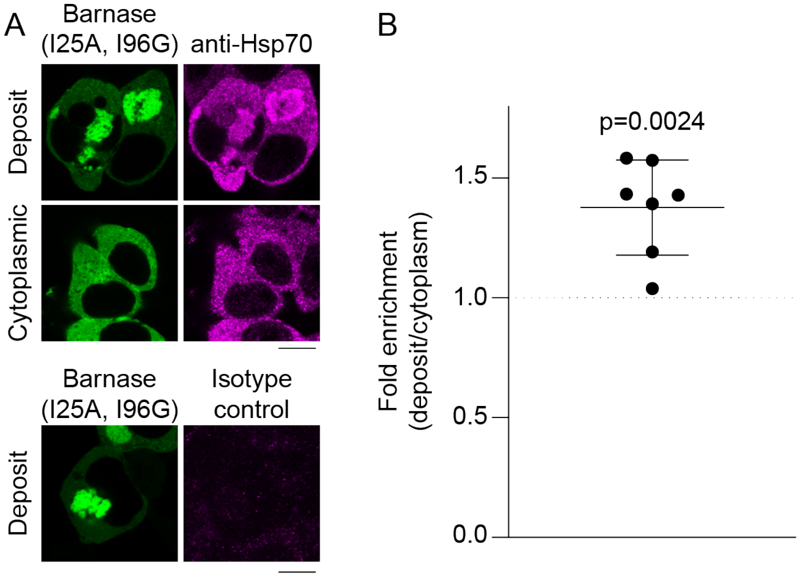


**Figure S4. Hsp70 binds barnase in both the dense and dilute phase. Related to Figure 5.** (A) Confocal images of Neuro2A cells transfected with the I25A, I96G barnase variant. Cells were stained by immunofluorescence for endogenous HSPA1A (Hsp70) or an isotype control as indicated. Anti-Hsp70 and isotype controls were imaged under the same conditions. Scale bar corresponds to 10 µm. (B) Graphs show quantitation of immunofluorescence. Mean and standard deviation shown, with p-value indicating results from a one-sample t-test.

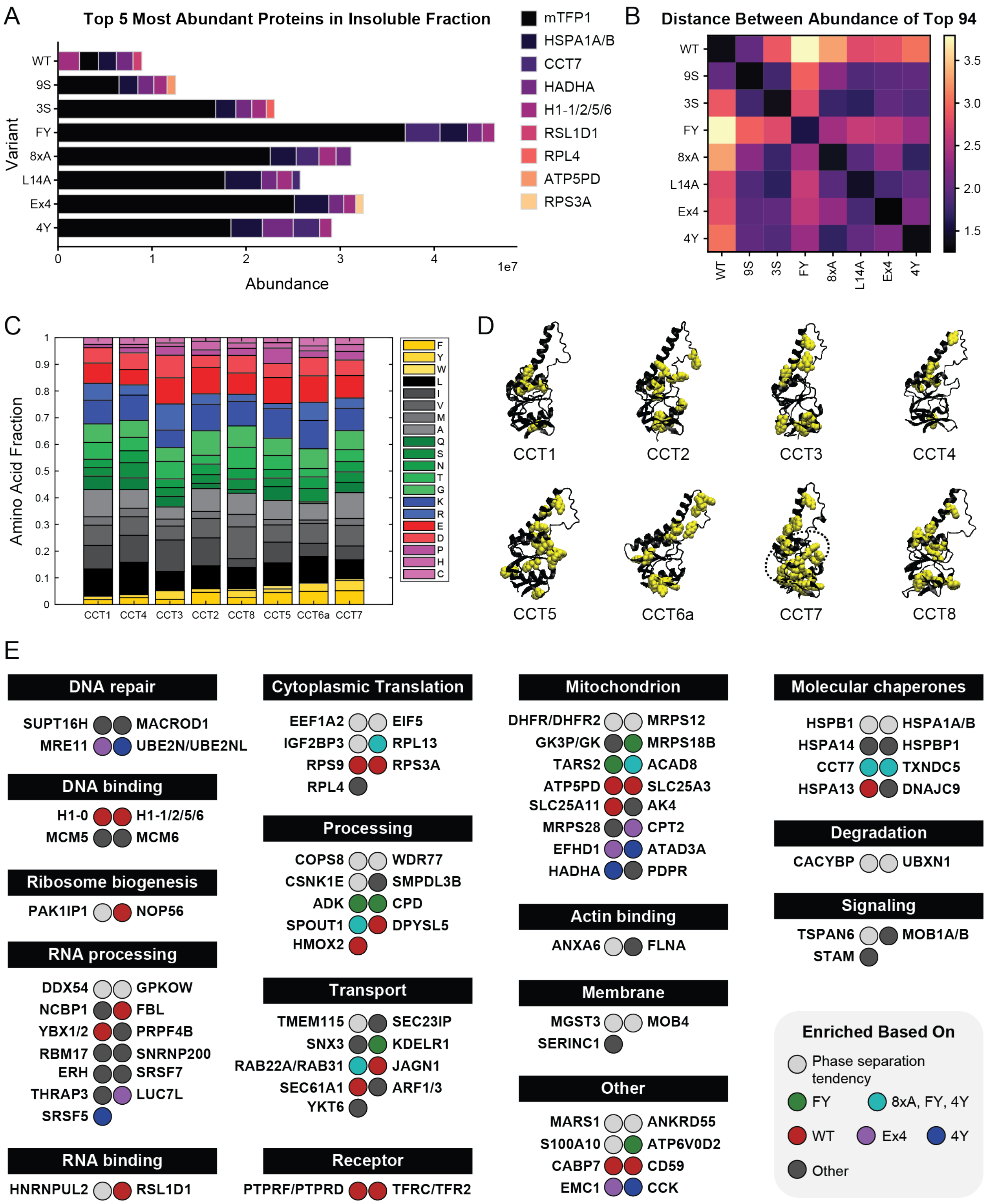


**Figure S5. Barnase UPODs show selectivity not based on overlapping or synergistic biological functions. Related to Figure 6.** (A) Abundance of the top 5 most abundant proteins in the insoluble fraction for each barnase variant within the set of the top 94 differently enriched endogenous proteins. Barnase (mTFP1) is the most abundant protein in each insoluble fraction, except for WT. (B) Mean Euclidean distance between $\log_{10}$(Abundance) vectors for each pair of barnase variants. Each vector includes the top 94 differently enriched endogenous proteins. Lower distances imply the compositional profiles of the insoluble fractions are more similar. (C) Amino acid fraction of the apical domain of each of the CCT subunits. (D) AlphaFold structures of the apical domains of each CCT subunit visualized using VMD (Humphrey et al., 1996; Jumper et al., 2021; Varadi et al., 2022). Aromatic residues are shown in yellow in space filling mode. Dashed oval denotes the substrate specificity region (Joachimiak et al., 2014). (E) Top 94 differentially enriched endogenous proteins grouped by function (Shemesh et al., 2021; Uhlen et al., 2015; UniProt, 2021). The circle color denotes how the protein is enriched in the variant specific insoluble fractions, i.e., the selectivity. No clear functional groups are observed based on selectivity.
